# Graph Machine Learning for Physiological Role Prediction in Protein Contact Networks: A Large-Scale Comparative Study on the Human Proteome

**DOI:** 10.64898/2026.04.10.717657

**Authors:** Mattia Cervellini, Alessio Martino

## Abstract

Proteins are fundamental macromolecules involved in virtually all biological processes. Their physiological roles are tightly linked to their three-dimensional structure, which can be naturally abstracted as Protein Contact Networks (PCNs), i.e., graphs where residues are nodes and edges encode spatial proximity. This representation enables the application of graph machine learning to address the protein functional annotation gap at proteome scale. In this work, protein function prediction is studied on the majority of the human proteome, focusing on enzymatic activity and enzyme class assignment as well-defined and biologically meaningful targets. A large-scale supervised analysis was conducted on PCNs derived from experimentally resolved human protein structures. Multiple graph-based learning paradigms were systematically compared under a unified evaluation protocol, including handcrafted graph embeddings, kernel methods, and end-to-end Graph Neural Networks (GNNs). Feature engineering approaches included (i) spectral density embeddings of the normalized graph Laplacian and (ii) higher-order topological representations based on simplicial complexes, with optional INDVAL-based feature selection. These representations were paired with linear, ensemble, and kernel classifiers, while GNNs were trained directly on raw PCNs exploiting a diverse set of message-passing strategies. Two tasks were considered: binary classification of enzymatic versus non-enzymatic proteins and multiclass prediction of first-level Enzyme Commission (EC) classes. Performance was assessed using repeated stratified splits to ensure robust and variance-aware evaluation. In the binary enzymatic classification task, the Jaccard-based graph kernel achieved the best performance with an adjusted balanced accuracy of 0.90, closely followed by GNNs trained end-to-end on PCNs. In the multiclass EC prediction task, GNNs demonstrated superior discriminative power, reaching an adjusted balanced accuracy of 0.92 and outperforming all explicit embedding and kernel-based approaches. Overall, results indicate that EC class prediction is intrinsically more complex than binary enzymatic discrimination and benefits from the higher expressivity of deep message-passing architectures. The findings demonstrate that graph-based representations of protein structure support competitive functional prediction at proteome scale, with classical kernel methods and modern GNNs offering complementary strengths in terms of accuracy, inference-time applicability, and flexibility.

## 1 Introduction

Proteins are essential macromolecules for cellular and organic physiology, underpinning many biological processes such as catalysing metabolic reactions, transducing and integrating biological signals, transporting ions and metabolites, providing mechanical scaffolding and regulating gene expression [2, 76]. Enzymes, in particular, accelerate chemical transformations by stabilizing transition states and lowering activation barriers, thereby enabling reaction rates compatible with life. Perturbations in protein function underlie diverse pathologies [67], from metabolic errors to cancer, so accurate functional annotation is a prerequisite for biological understanding and therapeutic discovery.

Despite decades of progress in biochemistry and structural biology, comprehensive functional characterization has not kept pace with the rapid accumulation of sequences and structures. Moreover, inferring function from sequence alone is complicated by factors like domain shuffling [34], convergent evolution [71], and the presence of multifunctional proteins [32]. Three-dimensional approaches on the other hand impose strict physicochemical constraints on activity: the geometry of active sites and the organization of co-factors collectively shape specificity and catalytic role. These considerations motivated the appearance of computational intelligence strategies that exploit structural information to propose functional hypotheses and guide experiments.

Within this context, inferring a protein’s physiological role from its structure is both timely and relevant. Structure-based learning approaches offer a faster complement to experimental annotation by leveraging patterns that recur across families and folds — such as common structural motifs and global architectural features — to distinguish protein functions. By using structural signals in predictive models, such methods aim to bridge the annotation gap and accelerate biological insight across large proteomes.

Proteins can be divided into two macro-categories: enzymatic and non-enzymatic. To carry out their function, enzymatic molecules bind substrates at key locations known as active sites which bind only specific substrates for specific reactions [36]. Each enzyme is associated with an Enzyme Commission (EC) Number [49], assigned by the Nomenclature Committee of the International Union of Biochemistry and Molecular Biology (IUBMB). The EC numbering system presents the hierarchical structure highlighted in Table 1.

**Table 1:** Hierarchy of the EC numbering system. Source: adapted from [70].

| Level | Description |
| --- | --- |
| EC x | **Main class** (e.g., Oxidoreductases, Transferases, etc.) |
| EC x.x | **Subclass** – type of compound or group involved |
| EC x.x.x | **Sub-subclass** – specific type of reaction |
| EC x.x.x.x | **Serial number** of the enzyme in its sub-subclass |

The IUBMB defined a total of seven distinct main classes of enzymes which are briefly described in Table 2 [30, 69].

**Table 2:**
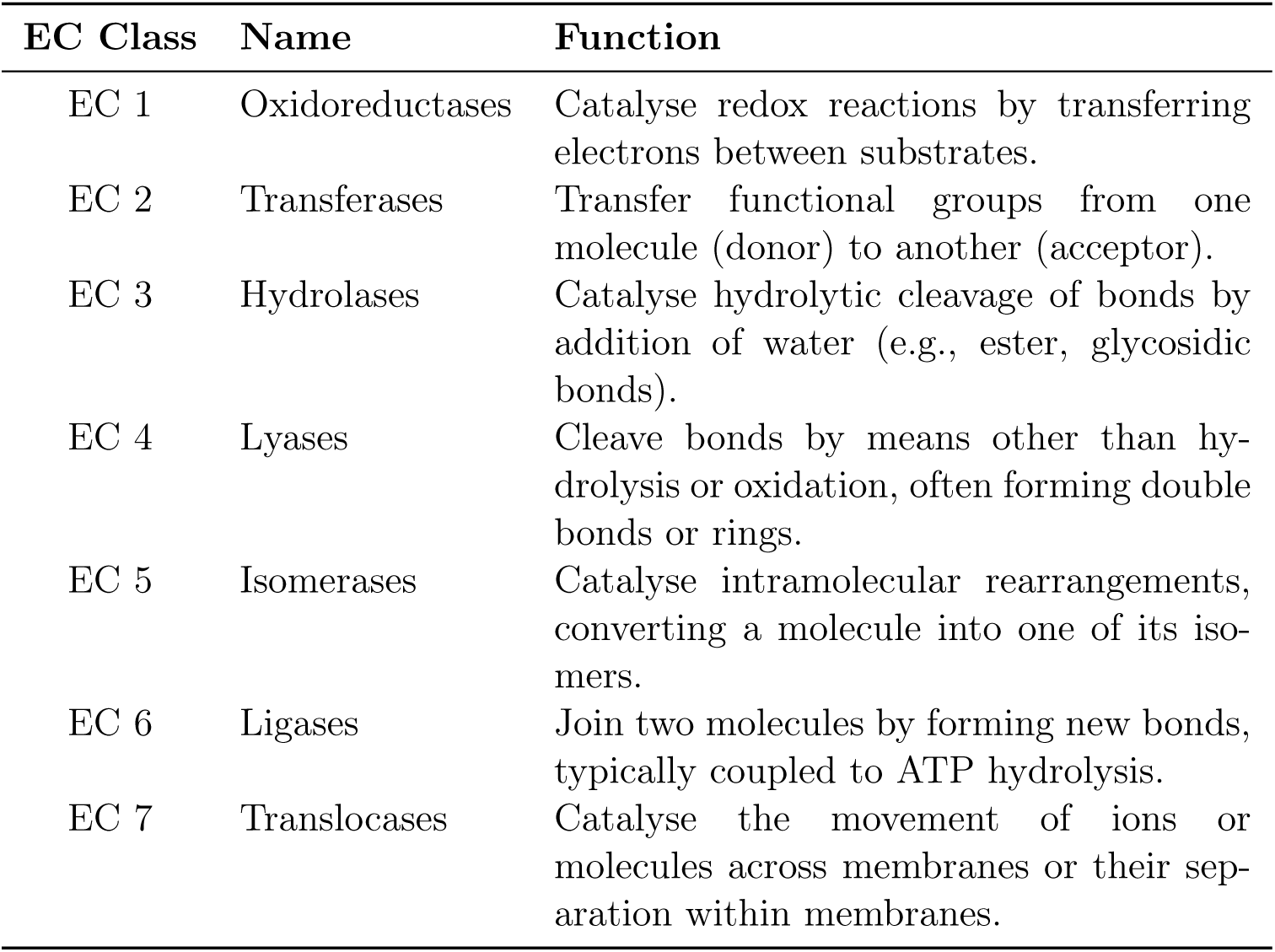
Main EC classes, names, and functions. Source: adapted from [45].

A powerful way to abstract the three-dimensional organization of proteins is to represent them as graphs. In this formulation, amino acid residues are mapped to nodes, while edges encode spatial proximity or chemical interactions between residues. The most common definition relies on a distance threshold applied to C*α* atoms resulting in a Protein Contact Net-work (PCN) [17]. This abstraction preserves essential information about the protein’s fold and intramolecular connectivity enabling systematic computational analysis.

Representing proteins as PCNs provides a natural substrate for exploring Graph Machine Learning (GML) methods, which are specifically designed to exploit relational and topological information of graph structures. Unlike conventional vector-based encodings, graph representations capture both local patterns and global architectural features. These properties make PCNs particularly suitable for function prediction tasks, where subtle structural characteristics often govern specificity and catalytic activity. Furthermore, graph abstractions are highly flexible: additional information such as residue type, chemical descriptors, or edge weights based on inter-residue distances can be seamlessly incorporated to enrich the representation.

By leveraging PCNs, modern learning algorithms can uncover structural signals fundamental for the analysis of complex topological structures like proteins. In this way, graph-based representations act as a bridge between raw structural data and predictive models, offering a framework for linking protein structure to physiological function.

This work investigates the capabilities of a diverse set of GML techniques, including embedding strategies inspired by spectral densities, algebraic topology, granular computing [6, 56], graph kernels, and Graph Neural Networks (GNNs), to recognize structural patterns for accurate protein classification. The focal objective is to classify proteins in the human proteome according to their physiological functions, formulated as two complementary tasks: (i) a binary discrimination between enzymatic and non-enzymatic proteins (hereinafter Task A), and (ii) a multiclass assignment of enzymatic proteins to their first-level EC classes (hereinafter Task B).

A distinctive contribution lies in the direct, uniform benchmarking of spectral- and algebraic topology-based descriptors against graph kernels and modern GNNs, an empirical comparison that appears not to have been systematically executed under shared experimental conditions in prior literature. The evaluation protocol is intentionally rigorous: stratified hold-out validation with fixed splits across all models, systematic hyperparameter optimization, and class-imbalance-aware performance measurements are employed to reflect real-world data characteristics. This design enables paired comparisons and strengthens the reliability of the conclusions. The analysis is conducted at proteome scale on ∼ 50, 000 unique human protein structures and spans 12 distinct combinations of learning algorithms and PCN representation techniques, ensuring a comprehensive and fair assessment of methodological strengths and limitations across all employed methodologies. These elements combined establish the work as a baseline reference for future studies, offering reproducible results and solid methodological guidelines to inform the design and evaluation of graph-based approaches for proteome-scale EC class annotation.

The rest of the work is organized according to the following structure: Section 2 reports an analysis of the previous scientific works with similar grounds; Section 3 describes the considered protein representations and Section 4 specifies the tested supervised learning models and performances evaluation criteria; Section 5 describes the data collection and provides a concise summary of the experimental pipeline; Section 6 highlights the empirical results deriving from the experiments; finally Section 7 presents a brief recap of the work and the final conclusions together with considerations regarding strengths, limitations and possibilities for future developments of the work.

## 2 Related Works

As anticipated in Section 1, in this work, the physiological role prediction is operationalised as two supervised tasks on protein structures. Methodologically, a two-stage formulation popularised in early structure-based work [8, 18, 19] is adopted: a binary classification between enzymatic and non-enzymatic proteins (Task A) followed by first level EC number classification on the enzymatic subset (Task B).

Plenty of studies have addressed protein classification by physiological role. An early example is [18], who aimed to distinguish enzymatic from non-enzymatic proteins without using sequence or structural alignments which were very popular at the time (e.g., BLAST [3] or FASTA [54]). They described proteins using both simple sequence-derived features (e.g., amino acid composition, etc.) and structure-derived features (e.g., secondary structure content, largest pocket size, etc.). A natural evolution of this study is [19] where the authors broadened the task to predicting also the first level EC number of the analysed proteins maintaining similar input features in a one-vs-one classification setting. Together, these studies established the feasibility of alignment-free, feature-based machine learning for physiological-role prediction while highlighting two themes that recur in later works: (i) structure-derived descriptors can add value beyond sequence-only baselines; and (ii) evaluation must account for class imbalance and protocol design.

Early work such as [8] modelled proteins at the level of secondary-structure elements, connecting them through sequential and structural edges. Subsequent studies [43] represented proteins as C*α* contact networks, defining edges between residues within a [4–8] Å distance range and summarizing the resulting graphs using topology- and spectrum-based descriptors combined with kernel methods and Support Vector Machine (SVM) variants; this line of work addressed first-level EC number prediction on a subset of the *E. coli* proteome. Related approaches [16] further integrated heterogeneous PCN- and sequence-derived descriptors into a dissimilarity space, evaluating multiple learning strategies including genetic feature selection with PARC/*ν*-SVM, isometric embeddings with standard classifiers, and cluster-based one-class models with learned feature weights.

Beyond physiological role prediction, PCNs have also been employed in a broader range of computational intelligence applications, including structural characterization, spectral analysis, and generative modelling [38,39,41]. This broader adoption reflects the versatility of PCNs as minimal representations of protein structure and motivates their use as the common graph substrate for the GML pipelines investigated in this work.

More recently, deep learning enabled structural representations to be learned directly from protein coordinates. An early example is EnzyNet [4], which represented enzymes as voxelised three-dimensional shapes and applied a 3D convolutional neural network for EC classification. Although this approach demonstrated that enzymatic classes can be inferred from spatial structure, it relied on a volumetric representation rather than on residue-level contact topology.

Among graph-based approaches, DeepFRI [25] represented proteins as C*α*–C*α* contact maps and used graph convolutions to predict EC and Gene Ontology annotations. Its proposed architecture combined the contact graph with residue-level embeddings extracted from a pretrained Long Short-Term Memory protein language model. More recent EC-focused methods have generally adopted richer structural or multimodal representations. GraphEC [65] constructs geometric graphs from ESMFold-predicted structures and combines them with protein language model embeddings, predicted active site information, and homology-based label diffusion. TopEC [72] instead employs localized three-dimensional graphs centred on candidate binding sites, incorporating distances and angles into the message-passing process.

Taken together, these studies illustrate a common tendency to enrich structural representations with pretrained sequence embeddings, explicit geometric quantities, active-site information, homology, or multimodal fusion. In contrast, the present work deliberately investigates a minimal PCN representation: an undirected, unweighted C*α* contact graph in which residue identity is the only node attribute, without sequence embeddings, distances, angles, edge attributes, or auxiliary prediction modules. To the best of our knowledge, previous works have not made this minimal representation the central subject of a proteome-scale controlled benchmark across both enzyme detection and first-level EC classification while comparing end-to-end GNNs with explicit topological embeddings and graph-kernel pipelines under identical experimental conditions.

## 3 Methods: Representation Strategies for Proteins

### 3.1 Proteins as PCNs

Protein structures were processed by creating PCNs following the same approach found in [16, 17, 43]. For each protein, starting from the 3D atomic coordinates of the amino acids, we considered their C*α* atoms as nodes in the resulting PCNs. C*α* atoms were connected if the Euclidean distance among them was in the range [4–8]Å, the lower bound was set in order to discard trivial first-neighbour interactions along the chain of a protein while the upper bound is set to approximately two van der Waals radii of C*α* atoms [53]. Outside such range residues are assumed to have no relevant interaction. The final PCNs are graphs whose nodes are labelled with the name of the residue corresponding to each C*α* atom. Edges were deliberately kept attribute-free to foster learning directly from the interactions of residues. Following the same rationale the resulting graphs have no notion of the original 3D space: they only retain information about the connectivity structure of the amino acids.

To provide an example, Figure 1 presents the structure of *Human Serum Albumin* both in its original 3D ribbon representation (Figure 1a) and in its PCN counterpart (Figure 1b) where each node is coloured according to the original residue of the relative C*α* atom. Note that nodes in PCNs were kept in the correct 3D coordinates for visualization purposes only as the analysis is entirely based exclusively on topological information.

**Figure 1:**
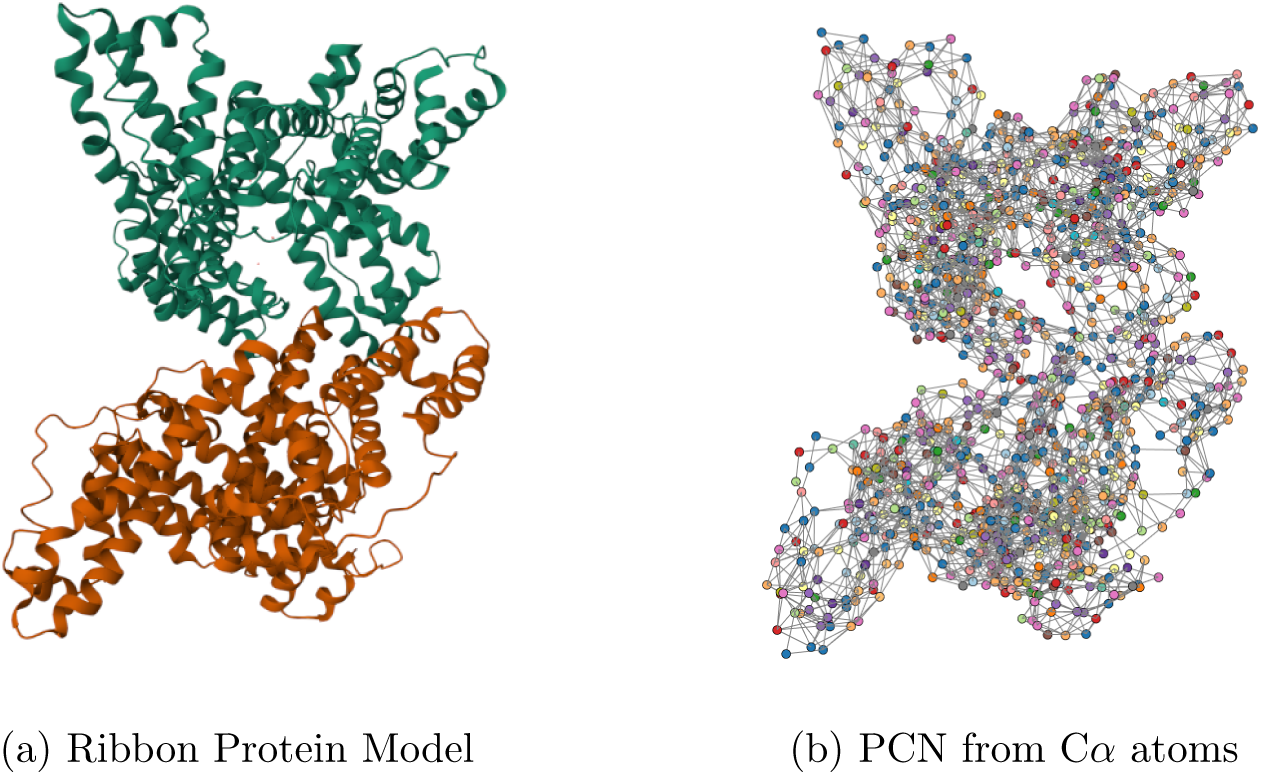
Representations for Human Serum Albumin (1AO6)

### 3.2 Graph Embedding via Simplicial Complexes

Simplicial complexes are a concept deriving from algebraic topology which has been widely explored in the domain of graph analysis [44, 52, 60]. A *k*-simplex can be seen as a convex hull formed by (*k* + 1) points of which each non-empty subset constitutes a face of such simplex which is itself a simplex of lower order. Building on these foundations and considering *s* a simplex of arbitrary order, a simplicial complex can be defined defined as a finite collection of simplices satisfying the following two properties:

1. if *s* ∈ S, every face in *s* is also included in S
2. if *s*_1_*, s*_2_ ∈ S, then *s*_1_ ∩ *s*_2_ is a face of both *s*_1_ and *s*_2_

By applying a simplicial complexes interpretation to PCN is it possible to create a *symbolic histogram* embedding of the protein structure by counting how many times each simplex appears in the protein [42]. This process however has limited representation capabilities when applied to out-of-the-box PCNs which are able to describe only pairwise relationships (PCNs are by definition simple, unweighted graphs).

To surpass this limitation, the edges of PCNs were aggregated creating clique hypergraphs. An hypergraph is a graph generalization in which edges (commonly known as hyperedges) can connect more than two nodes at the same time. Formally, an hypergraph can be defined as *H* = (ν*, ε*_h_) in which *V* is a finite set of nodes (same as plain graphs) while ε_h_ ∈ ν ^n^ is a set of hyper-edges which can connect an arbitrary number of nodes in simultaneously. Hypergraphs have been explored in literature to represent a great variety of complex systems (e.g., co-authorship networks, metabolic networks, brain functional networks, etc.) due to their flexibility and expressive power [12, 21, 27, 33, 47, 50, 75]. A clique hypergraph in particular is an hypergraph generated starting from a plain graph in which maximal cliques were substituted with hyperedges of the same order [5, 79]: a simple example of this process would be a triangle in the graph *G* described by edges (*A, B*), (*B, C*), (*A, C*) which gets projected in the hypergraph *H* as a single hyper-edge (*A, B, C*) . The same procedure is carried out for maximal cliques of all orders.

The instance matrix **X**^(S)^ resulting from such embedding via simplicial complexes has shape *n d* where *n* is the number of proteins in the dataset and *d* is the dictionary of distinct simplices in the dataset. Considering *c*(*H*_i_*, d*_j_) a function that counts how many times the simplex *d*_j_ appears in the clique hypergraph *H*_i_ the instance matrix **X**^(S)^ can be defined as

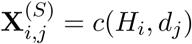

Given the two-step experimental framework of this work, two distinct instance matrices were constructed: (i) *X_A_*^(*S*)^ including a dictionary *d*_A_ of all simplices from both enzymatic and non-enzymatic proteins (∼ 16, 000 in total) for Task A; and (ii) *X_B_*^(*S*)^ including a dictionary *d*_B_ of only simplices coming from enzymatic proteins (∼ 13, 000 in total) for Task B.

#### 3.2.1 INDVAL Scores

Given the high dimensionality of the embedding presented in the previous section due to the proteome-scale dataset involved in the analysis, it was relevant to investigate whether it was possible to perform some degree of feature selection while maintaining stable classification performances. In order to carry out this procedure with a model-agnostic approach, the Indicator Value (INDVAL) score was investigated. The INDVAL score is a computational ecology derived indicator that accounts for both sensitivity and specificity and it was originally proposed to individuate the most characteristic species of a specific environment type [20]. According to the INDVAL criterion, a species *s* is a good representative for an environment type *E* if:

1. *s* is present only (or almost only) in environments *E* — proxy for specificity
2. *s* is present in all (or almost all) of environments *E* — proxy for sensitivity

The same rationale can be used for individuating signature substructures for a specific class of proteins. To adapt the INDVAL scores to the experimental framework of this work it is possible to draw a parallel where each simplex of node-labels in the aforementioned dictionary *d* is considered a species and each enzymatic class is considered as an environment type. It was therefore possible to construct a restricted version of the embedding presented in Section 3.2 where only the sub-structures with the best INDVAL scores are considered.

Considering *j* a specific physiological role and *i* a specific simplex, the INDVAL score can be defined starting from the following scores:

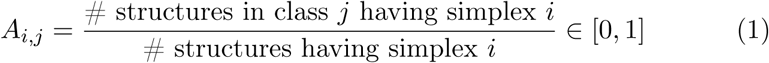

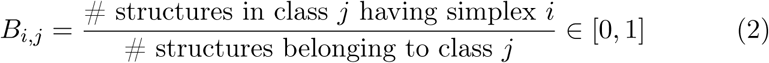

where Eq. (1) is a measure of specificity, maximized when simplex *i* appears only in class *j*; and Eq. (2) is a measure of sensitivity, maximized when all structures of class *j* have the simplex *i*. By combining Eqs. (1) and (2), the final INDVAL score *I*_i,j_ reads as:

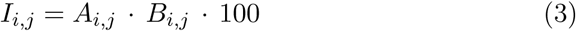

Given the bounds of Eqs. (1) and (2), it follows that *I*_i,j_ ∈ [0, 100]. This range admits the following interpretation: an INDVAL score *I*_i,j_ = 100 corresponds to a *perfect* simplex (i.e., substructure) from an information-theoretic perspective, meaning that observing the presence of this simplex is equivalent to knowing the class of the protein. In practical terms, simplex *i* would appear in *all* observations of class *j* and *never* in observations belonging to any other class.

Given a classification task where the symbolic histogram technique was used to embed the input data, it is possible to define its INDVAL matrix **I** of shape |*C| × |d|* where *d* is the dictionary of all the symbols in the dataset and *C* is the set of unique classes. Each value **I**_i,j_ corresponds to the INDVAL score of simplex *d*_j_ for class *C*_i_

Starting from **I** it is possible to select embedding features which had at least one of their INDVAL scores above a certain user-defined threshold *τ* . The choice of *τ* is not straightforward and strongly depends on the specific dataset as there is no guarantee of the existence of the *perfect* simplex: INDVAL scores in practice might be upper bounded at a value lower than 100. It is however possible to pick *τ* with a data driven approach thanks to some intuitive heuristics.

Considering the dual experimental framework of the paper, two distinct INDVAL matrices were defined: **I**_A_ with shape |*C*_A_| × |*d*_A_| for Task A and **I**_B_ with shape |*C*_B_| × |*d*_B_| for Task B. Figures 2 and 3 present the number of features (simplices) included in each dataset (Task A and B) for every possible *τ* . As expected the maximal INDVAL scores were far lower than 100 for both tasks, specifically they were 43.03 and 73.52 for Task A and B respectively. Notably most of the scores are near 0 as highlighted by power law shape of the distributions. The two vertical lines represent the two chosen thresholds *τ*_A_ = 6 and *τ*_B_ = 10 (for Task A and B respectively), such values were deemed appropriate in order to let the final INDVAL filtered embedding retain ∼ 10% of the original feature set while mostly respecting the elbow rule heuristic [68].

**Figure 2:**
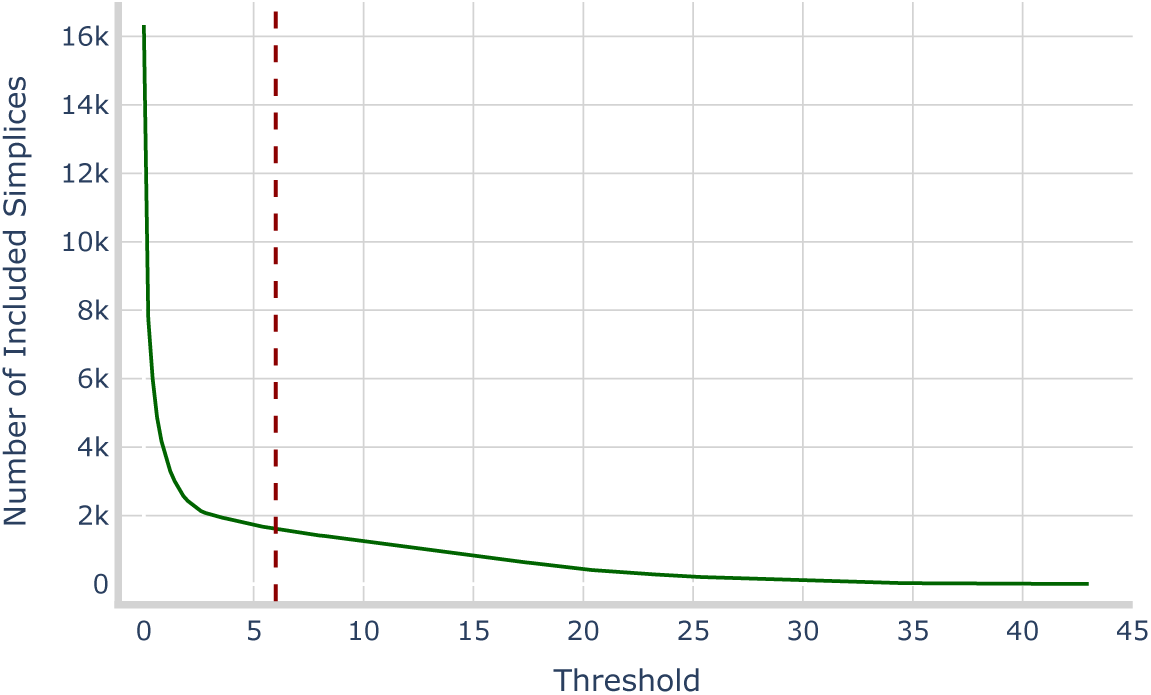
Number of features at various threshold levels for Task A.

**Figure 3:**
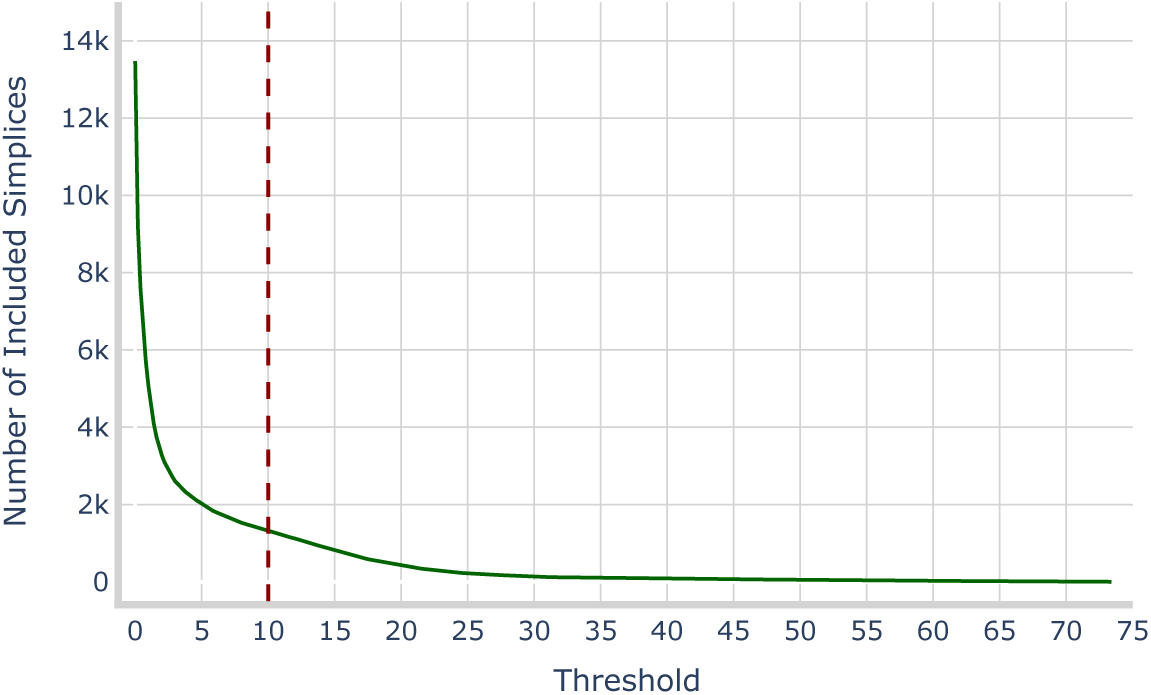
Number of features at various threshold levels for Task B.

The INDVAL feature selection procedure generated two reduced instance matrices: X*_A_*^(INDVAL)^ with ∼ 1, 600 features for Task A and X*_B_*^(INDVAL)^ with ∼ 1, 300 features for Task B. As a final remark, both X*_A_*^(INDVAL)^ and X*_B_*^(INDVAL)^ were obtained by filtering columns of X*_A_*^(*S*)^ and X*_B_*^(*S*)^ by comparing entries in **I**_A_ and **I**_B_ with *τ*_A_ and *τ*_B_ respectively.

### 3.3 Hypergraph Kernels

Graph kernels are among the most widely used techniques for GML tasks. The kernel methods presented in this section leverage the symbolic histogram embeddings described in Section 3.2 and are adapted from [44].

#### 3.3.1 Histogram Cosine Kernel

The Histogram Cosine Kernel (HCK) between any two PCNs can be computed as the cosine similarity between their respective symbolic histogram representations. Considering *x_i_* = X*_i_*^(*S*)^ the HCK between any two PCNs *i* and *j* can be defined as

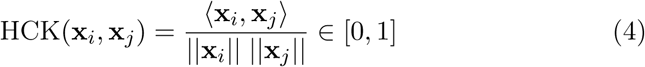

#### 3.3.2 Weighted Jaccard Kernel

The Weighted Jaccard Kernel (WJK) is computed as the ratio between the intersection and the union of two multi-sets. Considering again *d* as the dictionary of all simplices in the histograms and *x_i_* = X*_i_*^(*S*)^ the WJK between any two PCNs *i* and *j* can be defined as

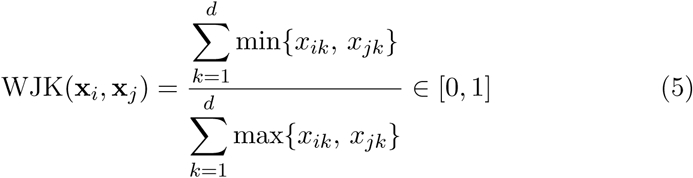

### 3.4 Graph Spectral Density

Let *G* = (*V, E*) be an undirected graph with *n* = |*V|* nodes, adjacency matrix **A**, and degree matrix **D** = diag(*D*_1_*, . . ., D*_n_), where *D*_i_ denotes the degree of node *i*. The graph Laplacian is defined as **L** = **D A** and plays a central role in spectral graph theory, as it encodes the connectivity structure of the graph [13, 24, 74]. A commonly used variant is the normalized Laplacian matrix, defined as

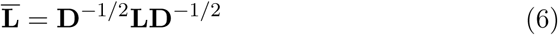

The normalized Laplacian has desirable theoretical properties: in particular it presents a spectrum that is invariant to d<u>eg</u>ree scaling and bounded within the interval [0, 2] [10]. The spectrum of **L^_^** (hereinafter referred to as graph spectrum) can be interpreted as a compact fingerprint regarding the connectivity properties of the graph itself. The cardinality of the graph spectrum however equals the number of nodes in the graph, which prevents a direct, size-agnostic use across different graphs which is key in supervised learning pipelines as the one presented in this work.

Thanks to the aforementioned properties of the graph spectrum it is possible to approximate the spectral density of a grap <u>h</u> via Gaussian Kernel Density Estimation (KDE) [41]. In order to do so, **L^_^** is treated as matrix with known spectral density *p*(*x*) [31, 43] that can be defined as

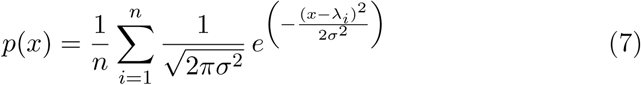

where *σ* represents the bandwidth of the kernel which, in order to make it graph-dependent, was set in accordance with the Scott’s rule [63]: considering *n* the number of nodes and *σ*^ the sample standard deviation of the eigenvalues, *σ* was defined as

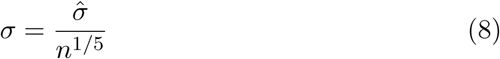

To move from the continuous density to a fixed-length feature vector, *B* = 200 uniformly spaced points in [0, 2] were sampled, producing a spectral embedding **s ∈** R^200^ for each PCN. This process yields comparable, size-agnostic descriptors that integrate seamlessly into the classification pipeline. As an example, Figure 4 shows the estimated spectral density for *Human Serum Albumin*; the corresponding embedding is given by the sequence of KDE evaluations at 200 grid points.

**Figure 4:**
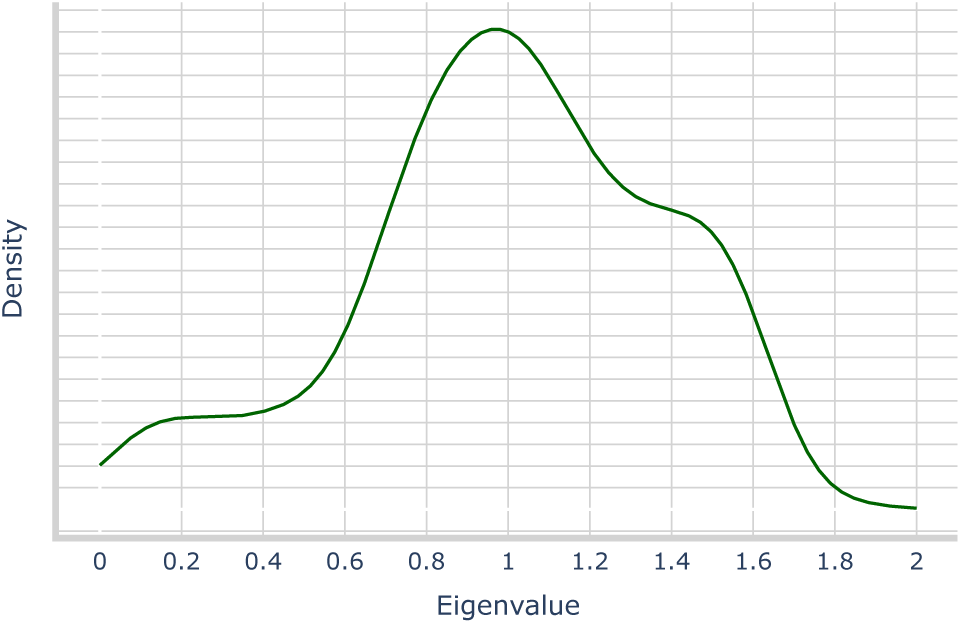
Spectral Density KDE Estimation for Human Serum Albumin

Applying the described spectral density embedding to each PCN yields two instance matrices, X*_A_*^(SPECTRAL)^ and X*_B_*^(SPECTRAL)^, corresponding to Task A and B, respectively. In both matrices, rows represent proteins and columns store the *B* = 200 evaluations of their spectral density on the uniform grid over [0, 2].

### 3.5 Summary of Representation Techniques

In summary, the experimental framework relies on three complementary representations derived from PCNs.

First, the simplicial complex embedding (Section 3.2) augments PCNs via clique hypergraphs to capture higher-order interactions; the resulting symbolic histograms offer direct compatibility with both standard machine learning algorithms and kernel methods emphasising label-aware local structures. The main limitations of said approach are its high dimensionality and the associated sparsity. Two non parametric hypergraph kernel approaches have been explored (Section 3.3) as they allow the convenient insertion of non-linear similarity metrics among graph structures. Kernels however do not produce explicit embeddings of the input data but yield complete Gram matrices which encode similarity among input objects.

Second, INDVAL-based feature selection (Section 3.2.1) retains substructures that are simultaneously specific to and prevalent within classes, yielding compact dictionaries (i.e., ∼ 10% of the original features considering the chosen thresholds) with a model-agnostic rationale. Threshold choice remains, however, data-dependent.

Third, the spectral density representation (Section 3.4) maps each PCN to a fixed-length vector by estimating the graph spectral density providing a size-agnostic summary of global connectivity that is robust to graph size; its weaknesses include loss of local motif identity and potential co-spectral graphs.

Taken together, the simplicial/INDVAL path prioritizes biologically in-terpretable, higher-order local structure, whereas spectral density contributes to a compact global fingerprint. Table 3 presents a schematic summary of the dimensions of each embedding methodology explored.

**Table 3:** Dimensionality of Embeddings.

| **Representation** | **Dimensionality** |  |
| --- | --- | --- |
|  | Task A | Task B |
| Simplicial Complexes | $\sim 16,000$ † | $\sim 13,000$ † |
| INDVAL | $\sim 1,600$ † | $\sim 1,300$ † |
| Spectral Density | 200 | 200 |

† Dimensionality varies slightly across splits of the data, refer to Section 4.4.

## 4 Methods: Machine Learning Pipeline

### 4.1 Tested Supervised Learning Approaches

The embedding strategies introduced in Sections 3.2, 3.2.1, 3.3, and 3.4, were evaluated using a set of established classification models selected after preliminary screening across multiple model families. The final choice balanced complementary inductive biases and practical considerations: (i) *ℓ*_1_-SVM [22, 78], chosen for computational efficiency and its embedded feature-selection behaviour that is well-suited to high-dimensional, sparse representations; (ii) kernel *ν*-SVM [61], selected for its flexibility in capturing nonlinear decision boundaries and, crucially, for its native support of kernelized learning, which makes it the natural classifier for the kernel representations introduced in Section 3.3; and (iii) a Random Forest [9], included as a robust, broadly competitive baseline that performs well across heterogeneous feature spaces. All three classifiers were applied to the vectorial embeddings (simplicial complexes, INDVAL, and spectral density). In contrast, for the kernel-based representations (HCK and WJK), only the kernel *ν*-SVM was considered, since it can directly operate on pre-computed Gram matrices. Table 4 presents a synthetic summary of which classification model was used in combination with each representation strategy.

**Table 4:** Summary of Representation Strategies and Associated Classifiers.

| Representation Strategy | $\ell_1$-SVM | ν-SVM | RF |
| --- | --- | --- | --- |
| Simplicial Complexes Embedding | ✓ | ✓ | ✓ |
| INDVAL Embedding | ✓ | ✓ | ✓ |
| Spectral Density Embedding | ✓ | ✓ | ✓ |
| Histogram Cosine Kernel | ✗ | ✓ | ✗ |
| Weighted Jaccard Kernel | ✗ | ✓ | ✗ |

### 4.2 GNNs

GNN architectures have been demonstrated to be extremely capable for graph classification often reaching state-of-the-art results. Figure 5 presents a conceptual summary of the core components of a GNN structure aimed at graph level machine learning tasks. All deep learning experiments of this paper were based on structures respecting the macro-phases highlighted in the figure: preprocessing, convolution, pooling, and finally classification.

**Figure 5:**
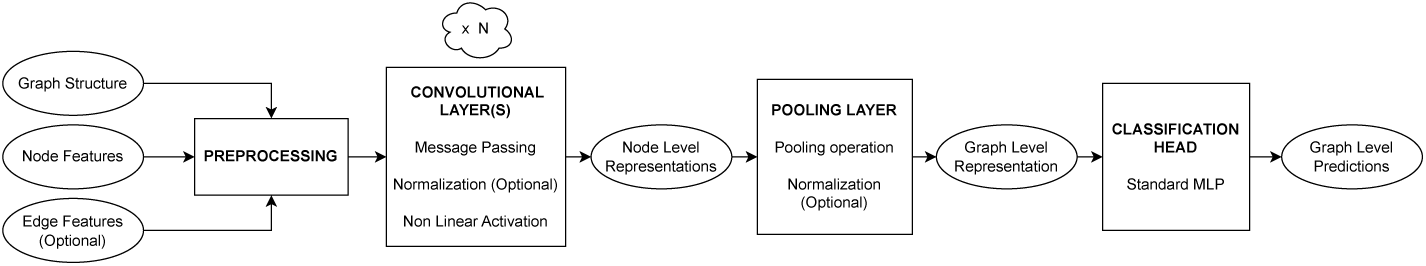
Sample GNN structure

In order to accommodate for the limited computing power available and to prevent known issues related to Message Passing (MP) such as over-smoothing (i.e., most node collapsing to near-identical representations after numerous MP steps [58]), the tested GNN architectures were limited to relatively shallow structures. Specifically all of the tested topologies had at most 5 MP steps, 5 classification head layers and latent dimensions (for each node representation) of at most 256. GNNs take as input PCN structures directly (via the associated incidence matrix) without the need for further processing or explicit embeddings. Once trained, the GNN can classify a newly constructed PCN through a single forward pass.

Recall that in this specific setting each node has one single categorical feature (i.e., its amino acid name), such label was represented either via One-Hot-Encoding (OHE) with a binary vector in 0, 1 ^21^ (20 dimensions for standard amino acids plus 1 for possibly unknown ones) or via dense learnable embeddings with a maximum of 128 dimensions.

In terms of MP strategies the exploration revolved around some of the most relevant MP paradigms in literature, namely Graph Convolution [48], SAmple and aggreGatE (SAGE) Convolution [29], Graph Convolutional Network (GCN) Convolution [35], Graph Isomorphism Network (GIN) Convolution [77], and Graph ATtention (GAT) Convolution [73]. Notably each explored GNN topology exploits only one strategy at a time with no change in the dimensions of node representations after any MP layer besides the first which projects node representations up to the chosen hidden dimensions of the network. As in Figure 5, MP was followed by graph pooling, which aggregates node representations into a single vector corresponding to a *graph latent representation*. Pooling was carried out either via standard permutation invariant aggregations (max/mean/sum) or via Attentional Aggregation [37]. After pooling the actual classification step was carried out thanks to standard MLP structure whose training included dropout to limit overfitting.

### 4.3 Performance Metrics

To provide a comprehensive evaluation of the proposed models, both easily interpretable performance metrics and metrics that are robust to class imbalance were considered. All the performance metrics employed in this work can be calculated starting from the standard entries of a canonical Confusion Matrix (CM), hereinafter abbreviated as TP for *True Positives*, TN for *True Negatives*, FP for *False Positives*, and FN for *False Negatives*. Note that the notion of CM also applies to multiclass classification, by expanding the matrix dimensions to match the number of distinct classes in the dataset. The metrics considered are the following:

- **Accuracy**: defined as

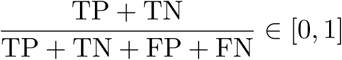

It represents the fraction of observations correctly classified by the model

- **Precision**: also known *Positive Predictive Value*, and defined as

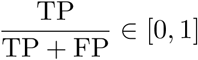

It represents the fraction of actually positive instances among the predicted positive instances.

- **Recall**: also defined as *Sensitivity*, and defined as

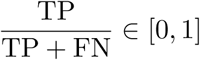

It represents the fraction of actually positive values correctly predicted by the model.

- **F1-Score**: defined as

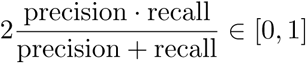

It is the harmonic mean of precision and recall and is often capable of conveying more information about model performance despite being harder to interpret.

- **Balanced Accuracy**: considering *C* the number of classes in the dataset Balanced Accuracy is defined as

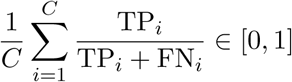

It represents the macro average of recall across all classes in the dataset and it avoids inflated performances in the case of imbalanced datasets which makes it often preferable to standard Accuracy. Balanced Accuracy can be adjusted for chance and is reformulated as:

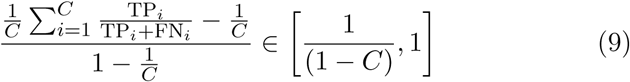

All presented performance measures can be easily extended to multiclass scenarios by averaging their binary version over all available classes. Adjusted Balanced Accuracy (ABA), specifically the formulation of Eq. (9), is naturally suited for multiclass scenarios and also avoids performance inflation on imbalanced datasets. For these reasons it was used as main performance metric throughout this work. Specifically, validation ABA was used as objective function value for the selection of the best set of hyper-parameters for all tested models (more details in the next Section).

### 4.4 Hyperparameter Optimization Strategy

Hyperparameter tuning was formalized as the global optimization of a black-box objective function *f* : *x* R over a (possibly) mixed (continuous, integer, categorical) search space *x*. The optimal set of parameters *x*^*^ can be defined as:

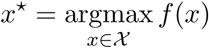

where each evaluation of *f* corresponds to training a model and evaluating its validation ABA under a specific hyperparameter configuration *x*. Hyperparameter search was implemented via multivariate Tree-Structured Parzen Estimators (TPE): a sequential model-based optimizer that replaces the usual Bayesian Optimization (BO) *p*(*y | x*) modelling [64] with a more convenient density estimation of *p*(*x | y*), yielding a sampling rule closely related to Expected Improvement while being naturally compatible with mixed and conditional search spaces [1, 7]. Its multivariate variant enables joint sampling and evaluation of subsets of hyperparameters, allowing dependencies between parameters to be exploited when sufficient joint observations are available.

In both experimental settings—Task A and Task B—the corresponding dataset was partitioned into three mutually exclusive, stratified training, validation, and test subsets, containing 60%, 20%, and 20% of the observations, respectively. This procedure was repeated using five independent split assignments. The same assignments were retained across all representation strategies and learning algorithms, allowing paired, like-for-like comparisons between pipelines.

For each split and pipeline, candidate configurations were trained exclusively on the training subset, while validation ABA was used as the objective function *f* (·) for hyperparameter optimization. All data-dependent processing steps, including feature scaling and simplex dictionary construction were performed without using the held-out test subset. After optimization, the hyperparameter configuration *x*^*^ achieving the highest validation ABA was retrained on the training subset and evaluated once on the corresponding test subset.

Tables 5–6 present a schematic summary of all hyperparameters that were tuned for each family of classifiers. Note that in both tables denotes conditional hyperparameters (e.g., dependent on kernel, architecture choices, etc.).

**Table 5:**
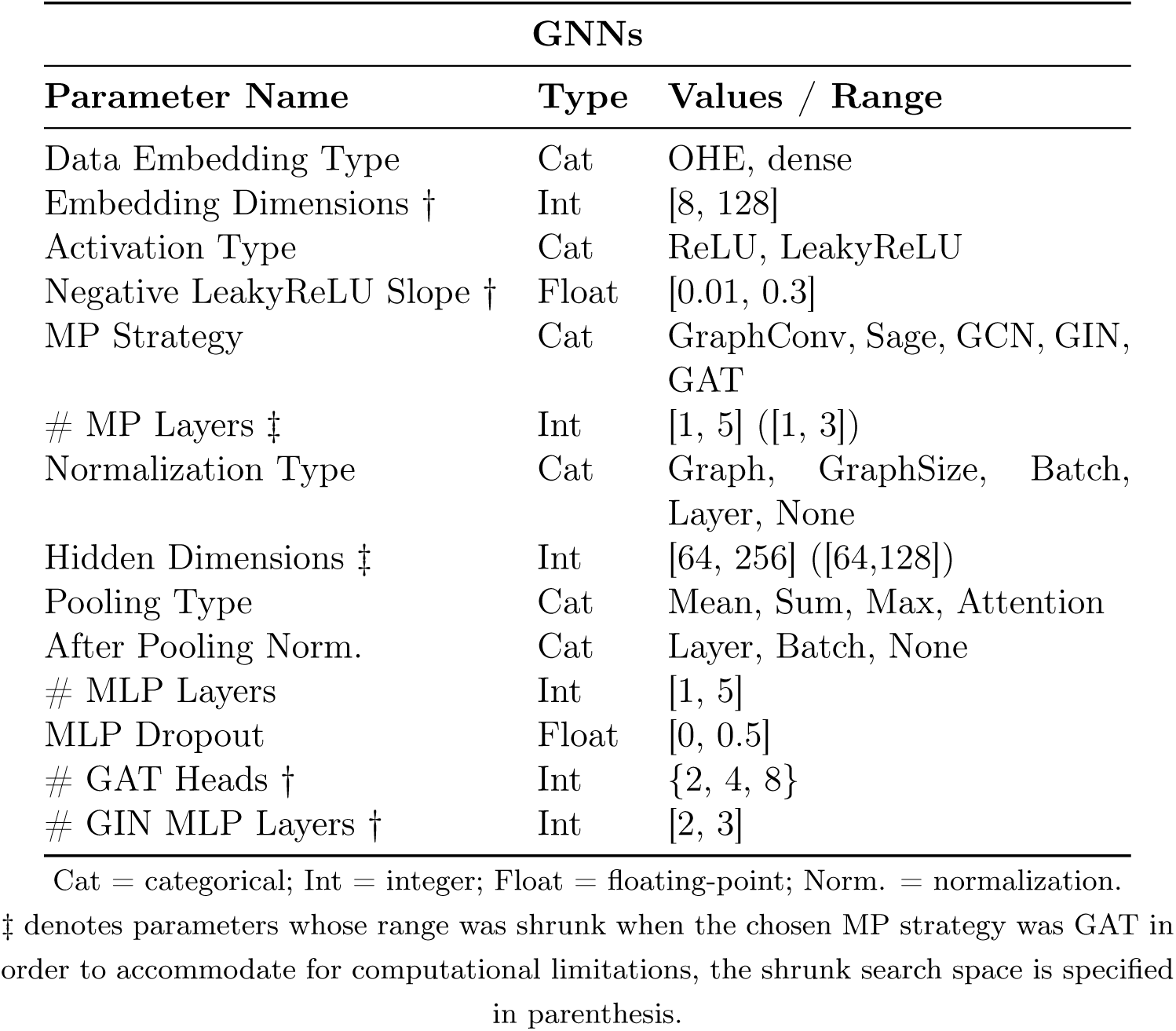
Search Spaces for GNN Model Family.

| GNNs |  |  |
| --- | --- | --- |
| Parameter Name | Type | Values / Range |
| Data Embedding Type | Cat | OHE, dense |
| Embedding Dimensions † | Int | [8, 128] |
| Activation Type | Cat | ReLU, LeakyReLU |
| Negative LeakyReLU Slope † | Float | [0.01, 0.3] |
| MP Strategy | Cat | GraphConv, Sage, GCN, GIN, GAT |
| # MP Layers ‡ | Int | [1, 5] ([1, 3]) |
| Normalization Type | Cat | Graph, GraphSize, Batch, Layer, None |
| Hidden Dimensions ‡ | Int | [64, 256] ([64,128]) |
| Pooling Type | Cat | Mean, Sum, Max, Attention |
| After Pooling Norm. | Cat | Layer, Batch, None |
| # MLP Layers | Int | [1, 5] |
| MLP Dropout | Float | [0, 0.5] |
| # GAT Heads † | Int | {2, 4, 8} |
| # GIN MLP Layers † | Int | [2, 3] |

Cat = categorical; Int = integer; Float = floating-point; Norm. = normalization.
‡ denotes parameters whose range was shrunk when the chosen MP strategy was GAT in order to accommodate for computational limitations, the shrunk search space is specified in parenthesis.

**Table 6:**
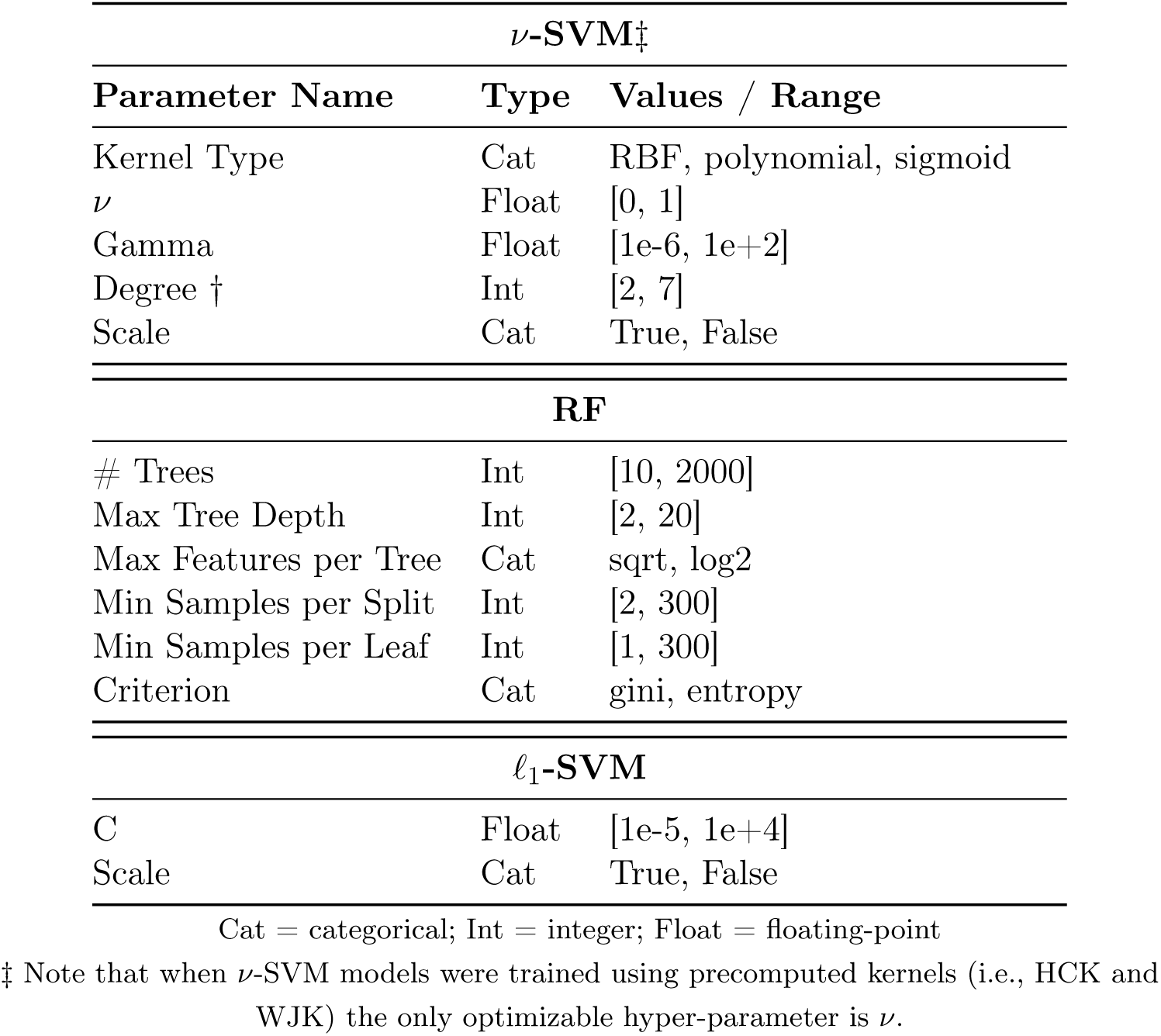
Search Spaces for Classical Model Families.

## 5 Data Collection and Experimental Setup

### 5.1 Data Collection and Dataset Construction

The complete data collection and PCN construction pipeline consisted of the following steps:

1. **Structure acquisition.** A total of 69, 979 human protein structures were downloaded from the Protein Data Bank^1^ (PDB) on March 1^st^, 2025.
2. **Structure parsing.** Each structure was processed to extract (i) the coordinates and residue identities of all C*α* atoms, (ii) the experimen-tal resolution, and (iii) the associated first-level EC annotation. When alternate atomic locations were present for the same residue their coordinates were averaged.
3. **Quality and annotation filtering.** Structures were discarded if they contained more than one first-level EC annotation, had missing experimental resolution or resolution exceeding 3 Å.
4. **Protein Contact Network construction.** Each retained structure was converted into an undirected, unweighted PCN. Each node *v* represents the C*α* atom of one amino-acid residue and is labelled according to its residue identity. An edge (*v*_i_*, v*_j_) was added when the Euclidean distance between the corresponding C*α* atoms was within [4, 8] Å, as detailed in Section 3.1. No edge attributes were assigned and the orig-inal three-dimensional coordinates were not retained. PCNs were discarded if they exhibited degenerate structures.
5. **Filtered dataset composition.** The two-staged filtering procedure (steps 3 and 4) yielded 48, 019 PCNs, including 26, 312 non-enzymatic and 21, 707 enzymatic proteins. The distribution of the enzymatic proteins across first-level EC classes is reported in Table 7.
6. **Task-specific dataset assembly.** Two datasets were derived from the filtered collection. Task A used all 48, 019 PCNs, labelled as enzymatic or non-enzymatic. Task B used the subset of 21, 679 enzymatic PCNs, each labelled according to its first-level EC class. Translocases (EC 7) was excluded from the dataset of Task B due to their limited prevalence. The two datasets were processed in parallel throughout all subsequent experimental stages.

**Table 7:** EC Number Distribution for Enzymatic Proteins.

| EC | Count | Prevalence |
| --- | --- | --- |
| 1 | 2959 | 13.63% |
| 2 | 8878 | 40.90% |
| 3 | 7181 | 33.08% |
| 4 | 1557 | 7.17% |
| 5 | 660 | 3.04% |
| 6 | 444 | 2.05% |
| 7 | 28 | 0.13% |

### 5.2 Experimental Setup

Figure 6 summarizes the experimental workflow adopted in this study for both Task A and Task B. Following data collection, filtering and PCN construction (Section 5.1), the resulting datasets were divided into five independent stratified training, validation and test splits, with identical assignments retained across all pipelines to ensure paired comparisons. The analysis then proceeded through two parallel branches: (i) a classical machine learning branch, combining explicit graph embeddings and kernel methods with suitable classifiers (Section 4.1), and (ii) an end-to-end GNN branch operating directly on raw PCNs and residue labels (Section 4.2). For each split and pipeline, hyperparameters were optimized to maximise validation ABA, after which the best configuration was retrained and evaluated once on the corresponding held-out test set (Sections 4.3–4.4). Finally, test metrics were aggregated across the five runs and the predictions of the various pipelines were compared.

**Figure 6:**
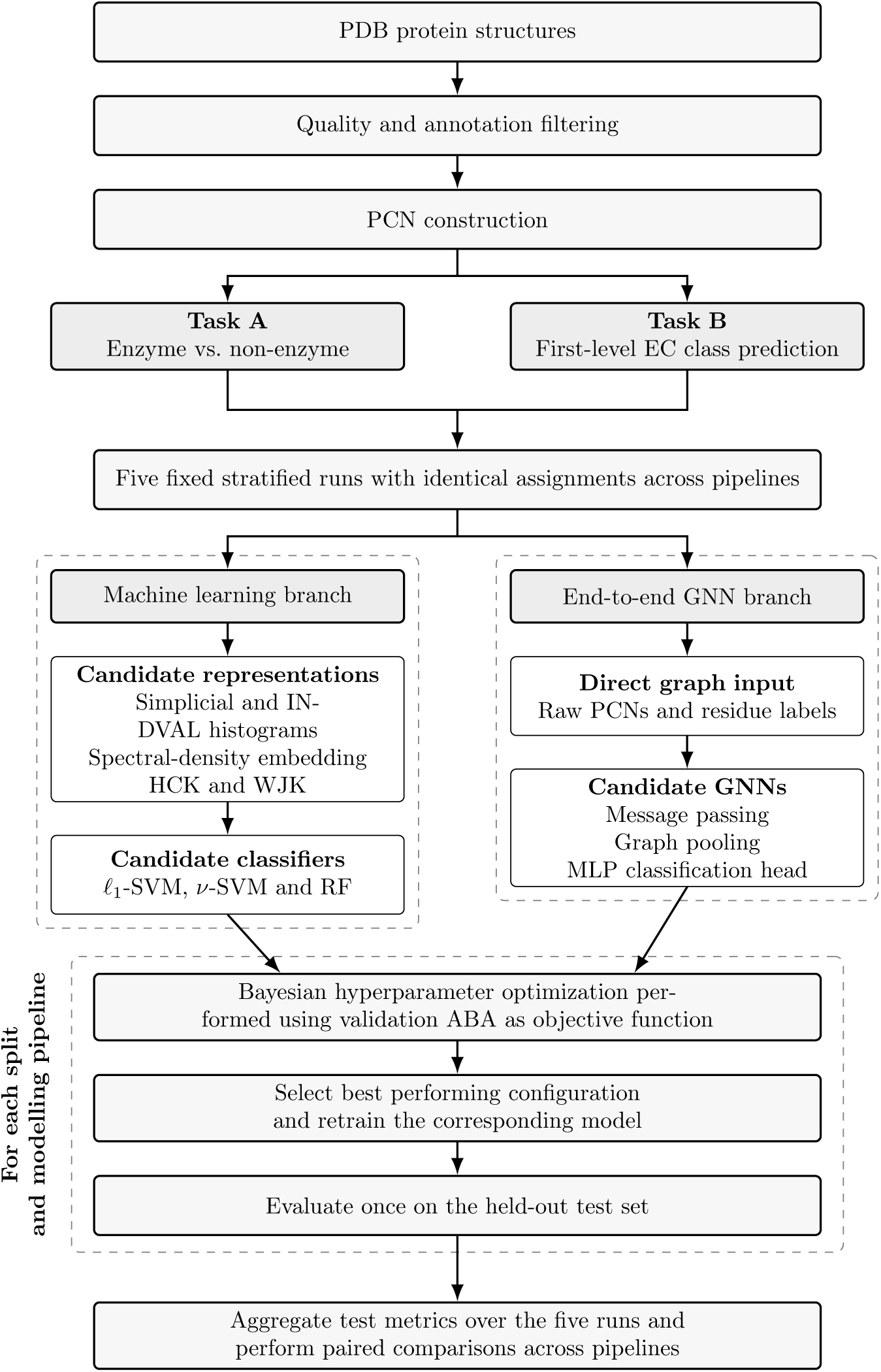
Unified experimental workflow of the paper.

To complement the aggregate performance measures with a formal statistical comparison, the McNemar’s test [46] was applied to the two models achieving the highest average test ABA within each task. The test was conducted separately for each of the five data splits using the paired predictions generated on the corresponding test set. This comparison was possible because identical test observations were used across all learning pipelines within each split. Statistical difference was assessed at *α* = 0.05 significance level.

The entire experimental pipeline was implemented in Python. Protein structures were parsed and processed using BioPython [14] and BioPandas [57]. PCN construction and other graph-related operations were performed using NetworkX [28]. Classical machine learning methods, data partitioning, and performance metrics were implemented using scikit-learn [55]. The GNN architectures were developed using PyTorch [51] and PyTorch Geometric [23]. Finally, hyperparameter optimization was conducted with Optuna [1].

## 6 Results

This section reports the performances of the tested methods together with the main outcomes of the analysis. The two experimental tasks are presented separately in Sections 6.1 and 6.2, respectively.

All tables in this section will present performance metrics averaged over the 5 distinct data splits (see Section 4.4) in the form avg ± std unless differently specified.

### 6.1 Task A: Enzyme vs. Non-enzyme Classification

#### 6.1.1 Simplicial Complexes Embedding

Table 8 highlights the performances on the test set for all models working with the PCNs simplicial complexes embedding for Task A. *ν*-SVM appears as the most powerful model among the tested ones, yet in this setting all classifiers perform quite similarly and *ν*-SVM takes the lead by a very small overall margin. *ℓ*_1_-SVM works especially well for this high dimensional embedding as its implicit feature selection allows for the sparsification of the solution, shortening running times and making predictions less noisy. Despite being a purely linear model *ℓ*_1_-SVM performs only 0.7% worse than *ν*-SVM and takes a fraction of the training time while simultaneously ignoring on average 78% of the input features in the process.

**Table 8:** Test set performances on simplicial complexes embedding – Task A.

| Model | ABA | ACC | F1 |
| --- | --- | --- | --- |
| ν-SVM | $0.874 \pm 0.003$ | $0.937 \pm 0.002$ | $0.931 \pm 0.002$ |
| $\ell_1$-SVM | $0.868 \pm 0.002$ | $0.934 \pm 0.001$ | $0.928 \pm 0.001$ |
| RF | $0.873 \pm 0.006$ | $0.940 \pm 0.003$ | $0.931 \pm 0.003$ |

RF models are inherently able to evaluate the importance of input features with the Mean Decrease in Impurity (MDI) method. *ℓ*_1_-SVM, on the other hand, are capable of deleting the least relevant features from the input data via intrinsic Lasso regularisation. These capabilities, combined with the fact that each feature in the simplicial complexes embedding represents the number of times a specific local sub-structure appears in a PCN, make it possible to analyse which sub-structures are more relevant when trying to separate Enzymatic and Non-Enzymatic proteins.

Figure 7a presents the top 10 most relevant features according to RF models. Raw RF feature importance scores were averaged across all 5 runs and normalized to 1. Figure 7b shows a similar plot on the hyperplane coefficients from the *ℓ*_1_-SVM: only features which were never removed in any of the five splits were considered, the absolute value of their coefficients was afterwards averaged across runs and normalized to 1 to facilitate comparison with Figure 7a.

**Figure 7:**
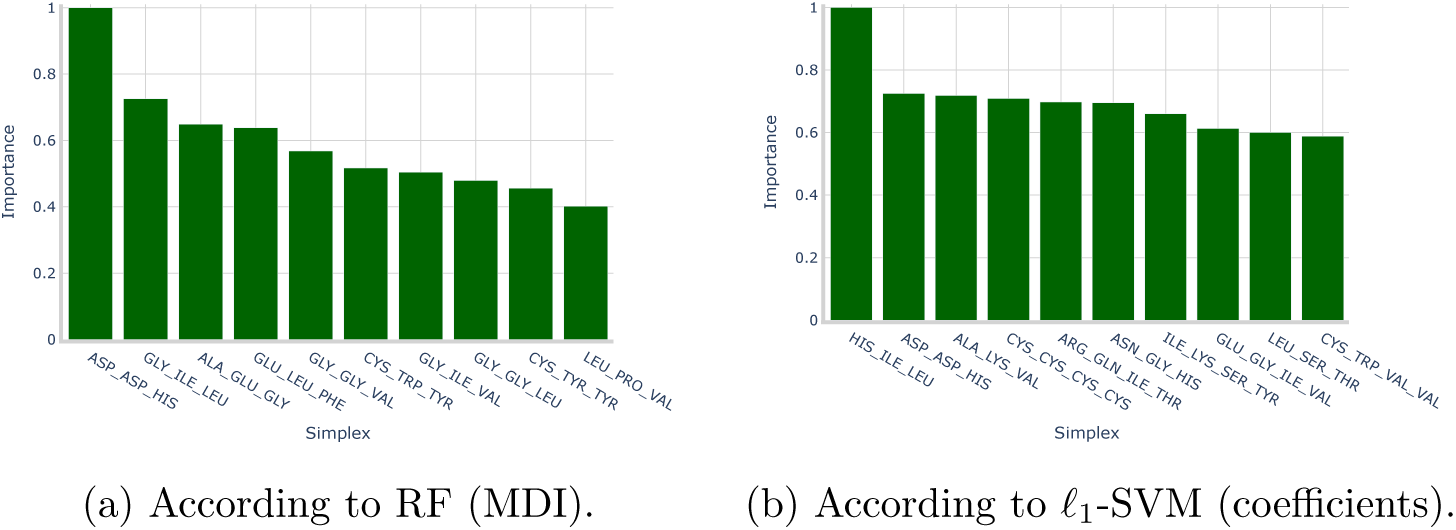
Top 10 most relevant simplices for Task A on the simplicial complexes embedding.

It is interesting to note how, despite RF and *ℓ*_1_-SVM being two very different models, the same simplex asp-asp-his appears among the most relevant ones for both models (more on this in Section 7).

#### 6.1.2 INDVAL Embedding

Table 9 shows the testing performances of all models when paired with the INDVAL embedding in Task A. In this setting, the RF model is the best performer with a test ABA 1.7% higher (on average) compared to *ν*-SVM. *ℓ*_1_-SVM underperforms slightly compared to the other methods: this is due to the feature vector being much smaller (∼ 10% of the simplicial complexes embedding) due to pre-filtering according to the INDVAL score. These conditions, combined with *ℓ*_1_-SVM algorithm pushing hyperplane coefficients to zero, was probably sub-optimal in this context and resulted in the removal of potentially relevant variables.

**Table 9:**
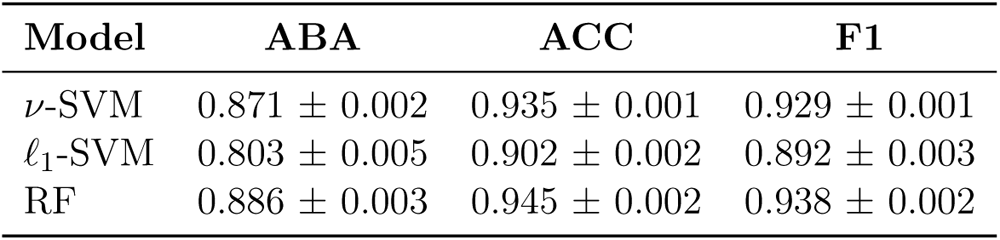
Test set performances on INDVAL-filtered embedding – Task A.

| Model | ABA | ACC | F1 |
| --- | --- | --- | --- |
| ν-SVM | $0.871 \pm 0.002$ | $0.935 \pm 0.001$ | $0.929 \pm 0.001$ |
| $\ell_1$-SVM | $0.803 \pm 0.005$ | $0.902 \pm 0.002$ | $0.892 \pm 0.003$ |
| RF | $0.886 \pm 0.003$ | $0.945 \pm 0.002$ | $0.938 \pm 0.002$ |

Alike the previous analysis, Figures 8a and 8b present the top 10 most relevant features for both RF and *ℓ*_1_-SVM. In this case there are no common simplices among the two models: however, it is interesting how the most important sub-structure according to RF is always asp-asp-his even after INDVAL feature selection. This phenomenon highlights how such 2-simplex is relevant both in terms of INDVAL score (as it surpasses the imposed threshold) and from a discriminative point of view (as it is consistently relevant across models).

**Figure 8:**
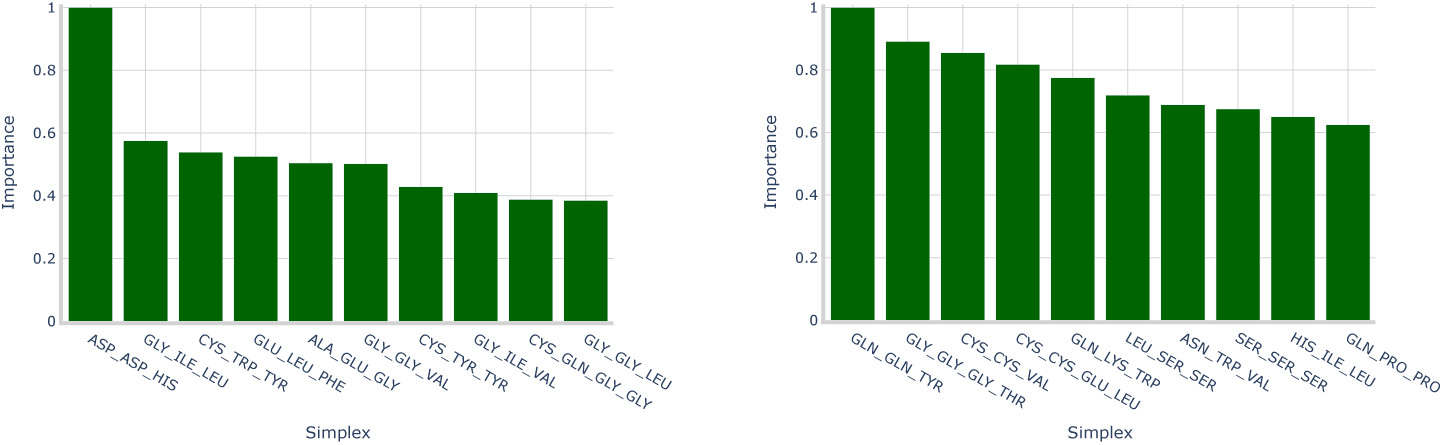
Top 10 most relevant simplices for Task A on the INDVAL-filtered embedding.

#### 6.1.3 Hypergraph Kernels

Table 10 shows *ν*-SVM testing performances on both kernel methods for Task A. Both kernels perform remarkably well, especially the WJK that exhibits the best results with a testing ABA of 0.900.

**Table 10:** Test set performances with kernel methods – Task A.

| Model | Kernel | ABA | ACC | F1 |
| --- | --- | --- | --- | --- |
| ν-SVM | HCK | $0.879 \pm 0.003$ | $0.939 \pm 0.002$ | $0.933 \pm 0.002$ |
| | WJK | $0.900 \pm 0.002$ | $0.951 \pm 0.001$ | $0.945 \pm 0.001$ |

The kernel-based approaches confirm the effectiveness of non-linear similarity measures between PCNs. It is however important to acknowledge that, compared to explicit embeddings, the computational limitations of kernel methods are relevant. For example the construction of full Gram matrices scales quadratically with the number of proteins and their interpretability is reduced compared to explicit embeddings where feature importances can be directly inspected.

#### 6.1.4 Spectral Density Embedding

Table 11 shows the testing performances for all models working with the spectral density embedding of PCNs in Task A.

**Table 11:** Test set performances on spectral embedding – Task A.

| Model | ABA | ACC | F1 |
| --- | --- | --- | --- |
| ν-SVM | $0.743 \pm 0.005$ | $0.872 \pm 0.003$ | $0.859 \pm 0.003$ |
| $\ell_1$-SVM | $0.351 \pm 0.006$ | $0.675 \pm 0.003$ | $0.655 \pm 0.004$ |
| RF | $0.719 \pm 0.007$ | $0.859 \pm 0.003$ | $0.848 \pm 0.004$ |

*ν*-SVM appears as the best performer with an average ABA of 0.743, 3.34% higher than RF. The *ℓ*_1_-SVM on the other hand exhibits substantially weaker performance: this behaviour can be largely attributed to the characteristics of the spectral density embedding. Since all graph spectra are supported in the range [0, 2], the KDE procedure samples exactly the same 200 evaluation points for every protein. This design ensures comparability across spectra but also introduces strong limitations. A prime example would be the Gaussian kernel smoothing enforcing continuity so, as a consequence, most estimated densities share very similar global shapes.

Under these conditions, KDE evaluations at two close points *x*_1_*, x*_2_ [0, 2] will yield nearly identical values. Hence, adjacent columns of the instance matrix are expected to be highly correlated. Figure 9 confirms this phenomenon by showing the correlation structure of **X**^(SPECTRAL)^: not only consecutive columns, but also columns near the spectral boundaries display high pairwise correlation. Being *ℓ*_1_-SVM by definition a linear classifier, the presence of strong linear correlation in the input features degrades its ability to identify sparse and discriminative subsets of variables.

**Figure 9:**
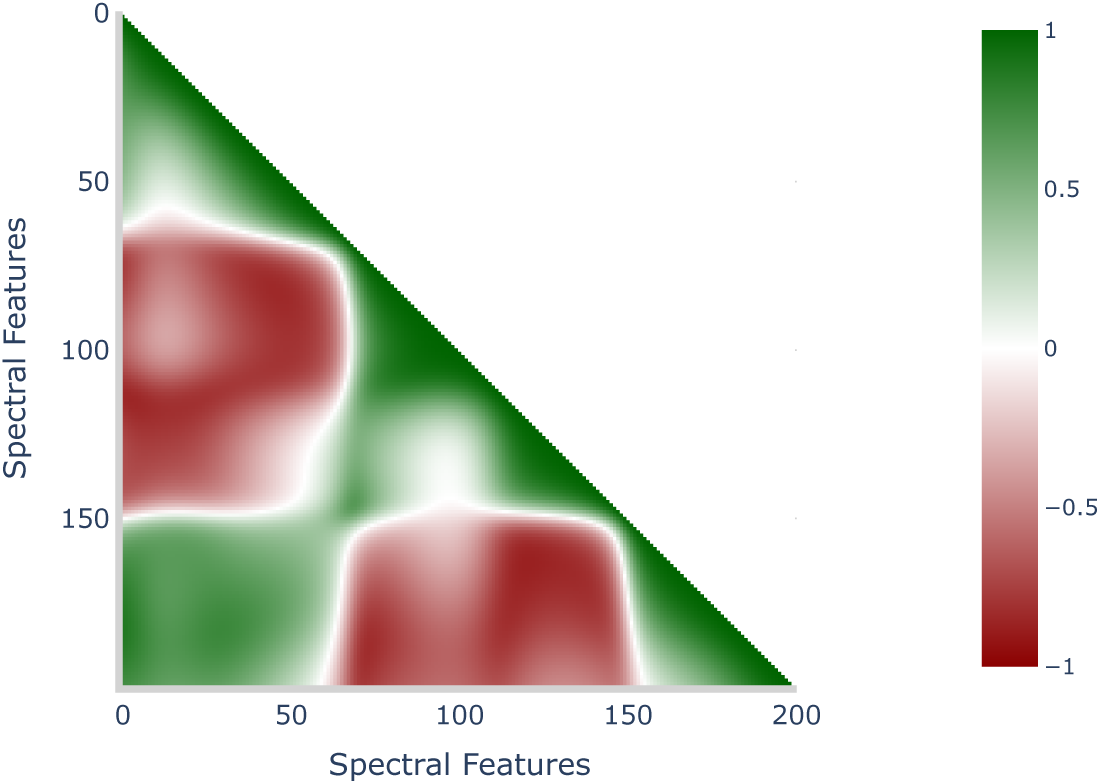
Pearson correlation coefficient of spectral features - Task A.

#### 6.1.5 Binary GNN

Table 12 presents GNN testing performances for Task A. From the 5 separate optimizations (see Section 4.4) some clear trends have emerged in the optimal structures. 4 out of 5 of the optimal topologies leveraged OHE for node labels and graph convolution for MP; at the same time, all configurations leveraged max pooling and batch normalization in MP layer. None of the optimal models leverages after pooling normalization and MLP heads have at most 3 layers. The hidden dimensions were relatively close to the upper bound of the associated search space, 3 out of 5 of the topologies selected 224 as optimal value while the others selected 192.

**Table 12:** GNN test set performances – Task A.

| Model | ABA | ACC | F1 |
| --- | --- | --- | --- |
| GNN | $0.898 \pm 0.005$ | $0.949 \pm 0.003$ | $0.944 \pm 0.003$ |

#### 6.1.6 Summary of Task A

Table 13 synthesizes the comparative results for the binary enzyme vs. non-enzyme classification task and three main conclusions emerge.

**Table 13:** Test set ABA for all models and representation strategies – Task A.

| Representation | ν-SVM | $\ell_1$-SVM | RF | GNN |
| --- | --- | --- | --- | --- |
| Spectral Density | $0.743 \pm 0.005$ | $0.351 \pm 0.006$ | $0.719 \pm 0.007$ | - |
| Simplicial Complexes | $0.874 \pm 0.003$ | $0.868 \pm 0.002$ | $0.873 \pm 0.006$ | - |
| INDVAL | $0.871 \pm 0.002$ | $0.803 \pm 0.005$ | $0.886 \pm 0.003$ | - |
| Histogram Kernel | $0.879 \pm 0.003$ | - | - | - |
| Weighted Jaccard Kernel | **$0.900 \pm 0.002$** | - | - | - |
| Raw PCNs | - | - | - | $0.898 \pm 0.005$ |

First, spectral density embeddings underperform across models (most markedly for *ℓ*_1_-SVM) due to the strong collinearity induced by KDE sampling similar distributions on a fixed grid.

Second, representations based on symbolic histograms of simplicial complexes, with or without INDVAL selection, are consistently strong and stable across models: RF and *ν*-SVM achieve competitive performance, and the linear *ℓ*_1_-SVM becomes viable in this high-dimensional, sparse regime thanks to its embedded feature selection.

Third, non-linear similarity functions on PCNs yield the best performances among all pipelines: the WJK attains the top average performance (test ABA of 0.900, slightly surpassing the HCK), albeit at the cost of quadratic Gram matrix construction and reduced interpretability compared to explicit embeddings. End-to-end deep learning on raw PCNs via GNNs performs almost on par with the best kernel approach (test ABA of 0.898), offering a compelling alternative that forgoes handcrafted features while preserving competitive generalization.

To determine whether the small difference between the two best-performing pipelines reflected a systematic difference in their predictions, the *ν*-SVM with WJK and the GNN were compared using McNemar’s test on each of the five test splits. A statistically significant difference was observed in two out of five splits, whereas the null hypothesis of equal error rates could not be rejected in the remaining three. Therefore, although the WJK pipeline achieved a marginally higher mean ABA, the statistical evidence for a consistent difference between the two approaches was limited across the five data splits.

Overall, Task A can be addressed effectively by both classical machine learning and deep learning approaches. When balancing performance and interpretability, simplicial complex-based embeddings offer an excellent trade-off; when maximizing absolute accuracy, the WJK is marginally superior; whereas GNNs provide a more direct inference pipeline, since a new PCN can be processed through a forward pass without the kernel evaluations required by kernel methods. Table 14 presents the test set confusion matrix of the *ν*-SVM model on the WJK as a final intuitive visualization of the maximal performances attained in Task A.

**Table 14:** ***ν*-SVM with Jaccard Kernel confusion matrix – Task A.**

|  |  | Predicted |  |
| --- | --- | --- | --- |
|  |  | Non-Enzyme | Enzyme |
| True | Non-Enzyme | 5046 | 216 |
|  | Enzyme | 246 | 4095 |

### 6.2 Task B: Multiclass Physiological Role Prediction

#### 6.2.1 Simplicial Complexes Embedding

Table 15 shows the testing performances of all models working with the simplicial complexes embedding in Task B. Such results highlight the effectiveness of linear models in high-dimensional and sparse feature spaces: *ℓ*_1_-SVM consistently emerges as the best performing classifier, surpassing both nonlinear approaches such as *ν*-SVM and more flexible ensemble methods like RFs. This outcome shows off how the implicit feature selection mechanism induced by the *ℓ*_1_ regularization is highly effective in this context. The number of simplices deemed as irrelevant by *ℓ*_1_-SVM averages to 10, 600 out of the 13, 000 in the dataset which means that approximately 81% of the features were ignored for classification purposes.

**Table 15:** Test set performances on simplicial complexes embedding – Task B.

| Model | ABA | ACC | F1 |
| --- | --- | --- | --- |
| ν-SVM | $0.864 \pm 0.009$ | $0.949 \pm 0.003$ | $0.948 \pm 0.003$ |
| $\ell_1$-SVM | $0.902 \pm 0.013$ | $0.953 \pm 0.006$ | $0.954 \pm 0.006$ |
| RF | $0.882 \pm 0.009$ | $0.946 \pm 0.005$ | $0.946 \pm 0.005$ |

From a feature importance perspective, the same methodology adopted for Task A can be applied also for Task B to identify the most relevant substructures within the simplicial complexes embedding. Figure 10 presents the 10 most relevant simplices in the dataset according to the RF model, it is interesting to see again how the 2-simplex asp-asp-his consistently emerges as the most influential feature across both classification scenarios, reinforcing its role as a potentially critical structural motif for enzymatic function recognition.

**Figure 10:**
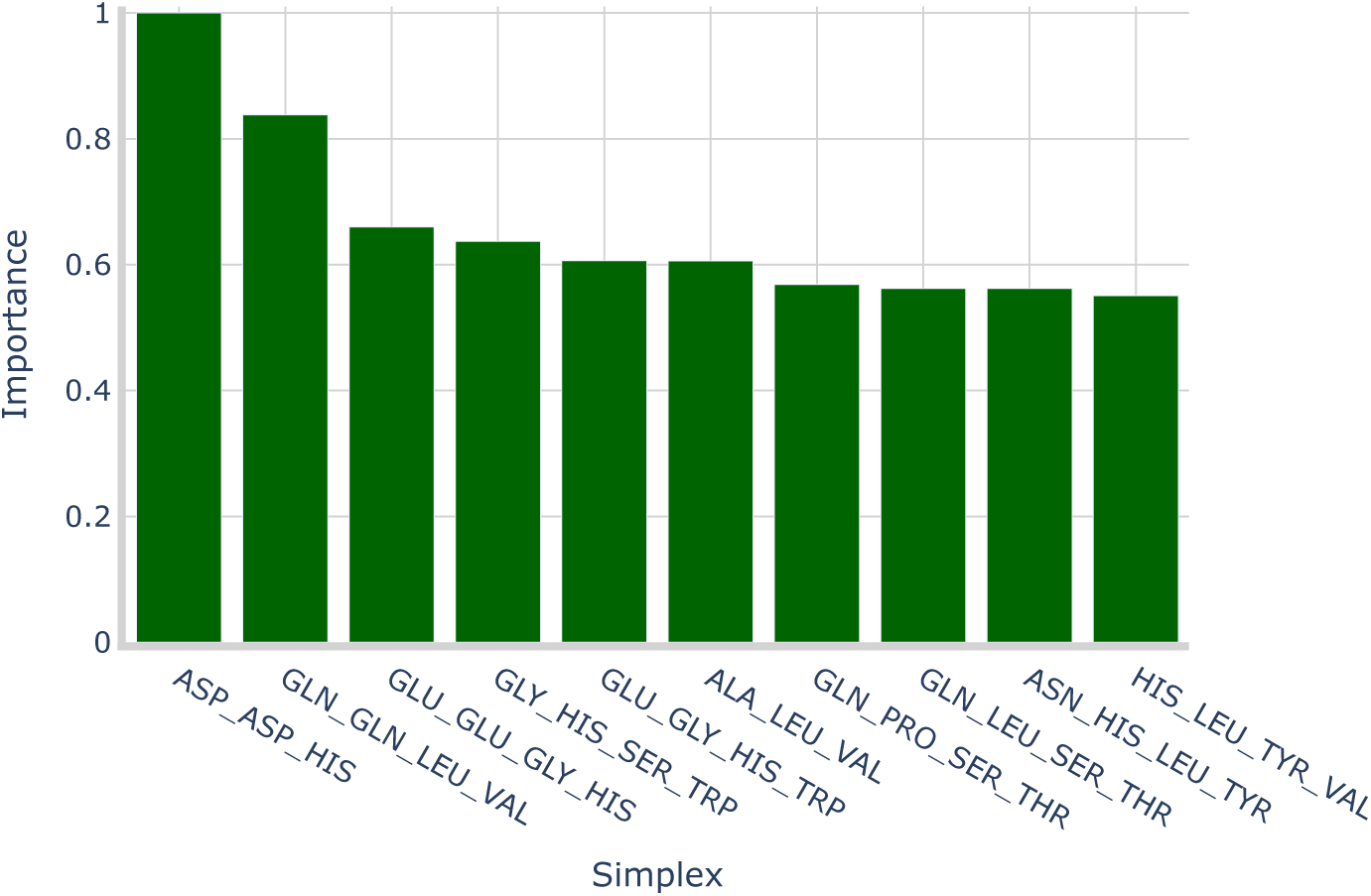
Top 10 most relevant simplices according to RF (MDI) on the simplicial complexes embedding – Task B.

When it comes to *ℓ*_1_-SVM the feature importance analysis for a multiclass scenario becomes less intuitive: in accordance with the One-vs-Rest (OvR) strategy, a different classifier is built for each class in the dataset and each of the classifiers is a completely independent model. At inference time all the models make their predictions and the one having the strongest prediction is the one effectively assigning the label.

Figure 11 shows feature importance plots similar to Figure 7b for each of the 6 distinct EC Classes in the dataset for Task B. These per-class coefficients must be interpreted within the OvR setting: each panel reflects a classifier trained against all remaining classes, hence coefficients encode class-specific discriminants and are not directly comparable across classes nor to RF MDI scores. The patterns highlighted by *ℓ*_1_-SVM complement those highlighted by RF indicating both shared and class-dependent structural signals captured by models with different classification strategies. Since each OvR classifier is fit independently, raw coefficient magnitudes reflect its own regularization path and are best read as within-class rankings rather than absolute importances.

**Figure 11:**
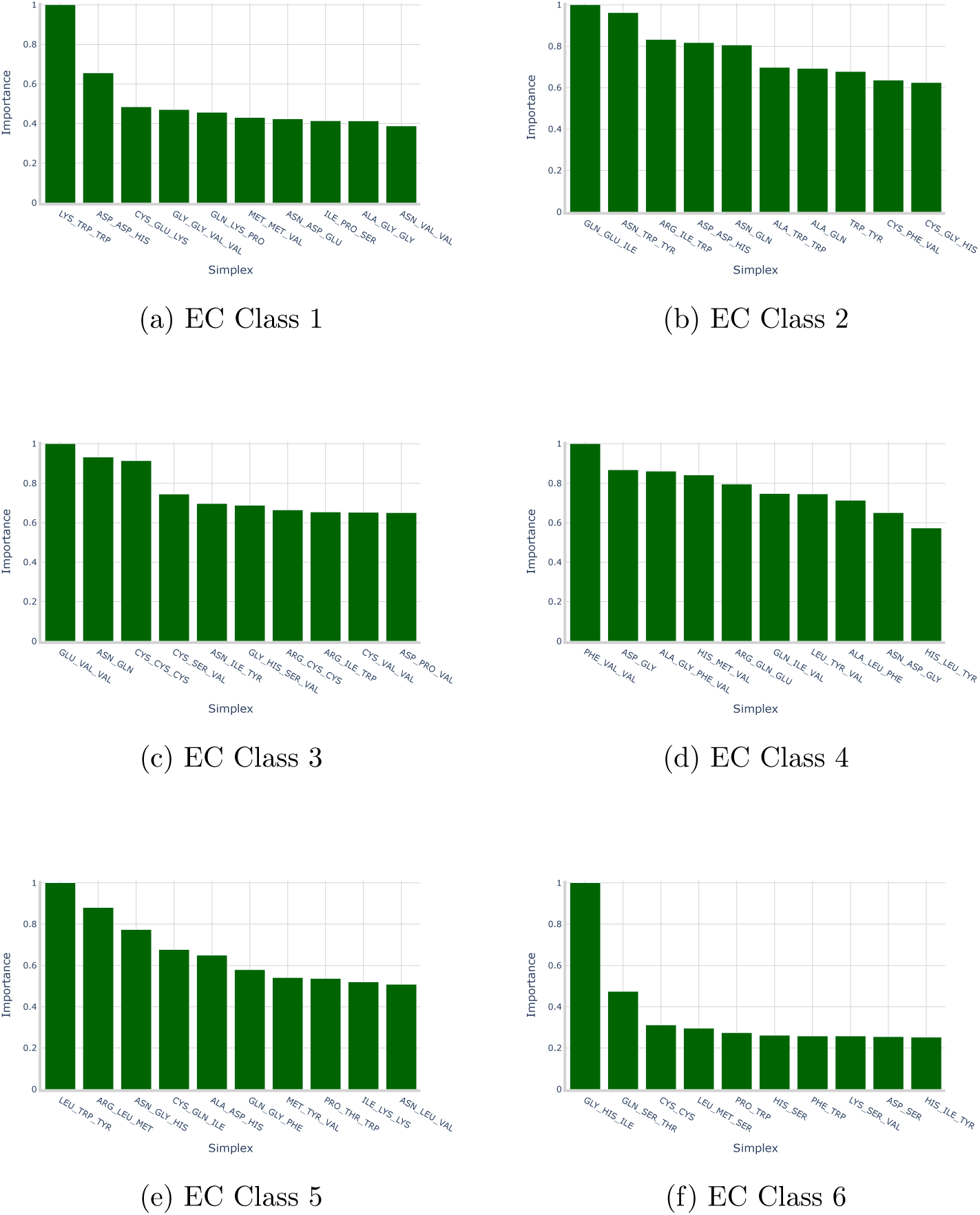
Top 10 most relevant simplices for each EC class according to *ℓ*_1_-SVM (coefficients) on the simplicial complexes embedding – Task B.

Despite such limitations it is worth noting that the 2-simplex asp-asp-his appears of great importance for both Oxidoreductases (EC Class 1) and Transferases (EC Class 2) as well as the notable differences in the distributions of the top 10 coefficients: for classes 2 to 5 the relevance distribution appears fairly evenly distributed (i.e., many of the best simplices have relatively close importance scores). Classes 1 and 6 on the other hand exhibit the opposite pattern, only the best 2 simplices have relatively similar importances while others display a great drop in relevance.

A final consideration on *ℓ*_1_-SVM implicit feature selection is that there is no single simplex that has been selected as relevant for all classes, this signals that different enzymatic classes are effectively differentiated by diverse structural motifs which further strengthens the grounds for PCN embedding via simplicial complexes.

#### 6.2.2 INDVAL Embedding

Table 16 presents the testing performances of all models working with the INDVAL embedding in Task B. Also in this case the best performing model in terms of ABA is *ℓ*_1_-SVM, although ranking the worst in terms of standard accuracy. It is worth recalling that standard accuracy measures the proportion of correctly classified samples across the entire dataset, it is therefore strongly biased towards the most represented classes. ABA on the other hand assigns the same weight to each class regardless of its frequency. As a consequence, a higher ABA accompanied by a lower accuracy indicates that the classifier is able to capture informative patterns also in minority classes even if this comes at the cost of slightly reduced performance on majority ones. In practice, such behaviour suggests that the model is better balanced across the class spectrum and performs well on under-represented EC classes. The strong *ℓ*_1_ penalty allows for a highly regularized model which on the INDVAL dataset excludes on average ∼ 592 features from the classification, corresponding to ∼ 46% of the entire feature set. As expected, the feature selection is less harsh compared to the previous section due to features being already pre-selected according to the INDVAL criterion. It is also important to note how, compared the results in Table 15 the INDVAL embedding achieves an average testing ABA only 1% lower than the full simplicial complexes embedding while being less than 10% the size. With this in mind the INDVAL scores appear to be exceptional in selecting the most relevant sub-structures for correct functional annotation.

**Table 16:** Test set performances on INDVAL-filtered embedding – Task B.

| Model | ABA | ACC | F1 |
| --- | --- | --- | --- |
| ν-SVM | $0.861 \pm 0.008$ | $0.946 \pm 0.003$ | $0.945 \pm 0.003$ |
| $\ell_1$-SVM | $0.893 \pm 0.012$ | $0.942 \pm 0.006$ | $0.943 \pm 0.005$ |
| RF | $0.888 \pm 0.010$ | $0.948 \pm 0.006$ | $0.948 \pm 0.006$ |

Figure 12 shows the feature importance plot for the RF model which again indicates as most important feature the 2-simplex asp-asp-his con-firming even further the relevance of such substructure. Figure 13 on the other hand presents the same visualization as in Figure 11 but for the INDVAL embedding. It is easy to see how the feature importances are more uniformly distributed compared to the ones coming from the full simplicial complexes embedding (see Figure 11) due to the feature selection carried out via INDVAL scores that allows the model to consider only a very small class-relevant subset of features whose relevance is smoothed due to the decreased dimensionality of the dataset. The 2-simplex asp-asp-his appears also here for both EC classes 1 and 2.

**Figure 12:**
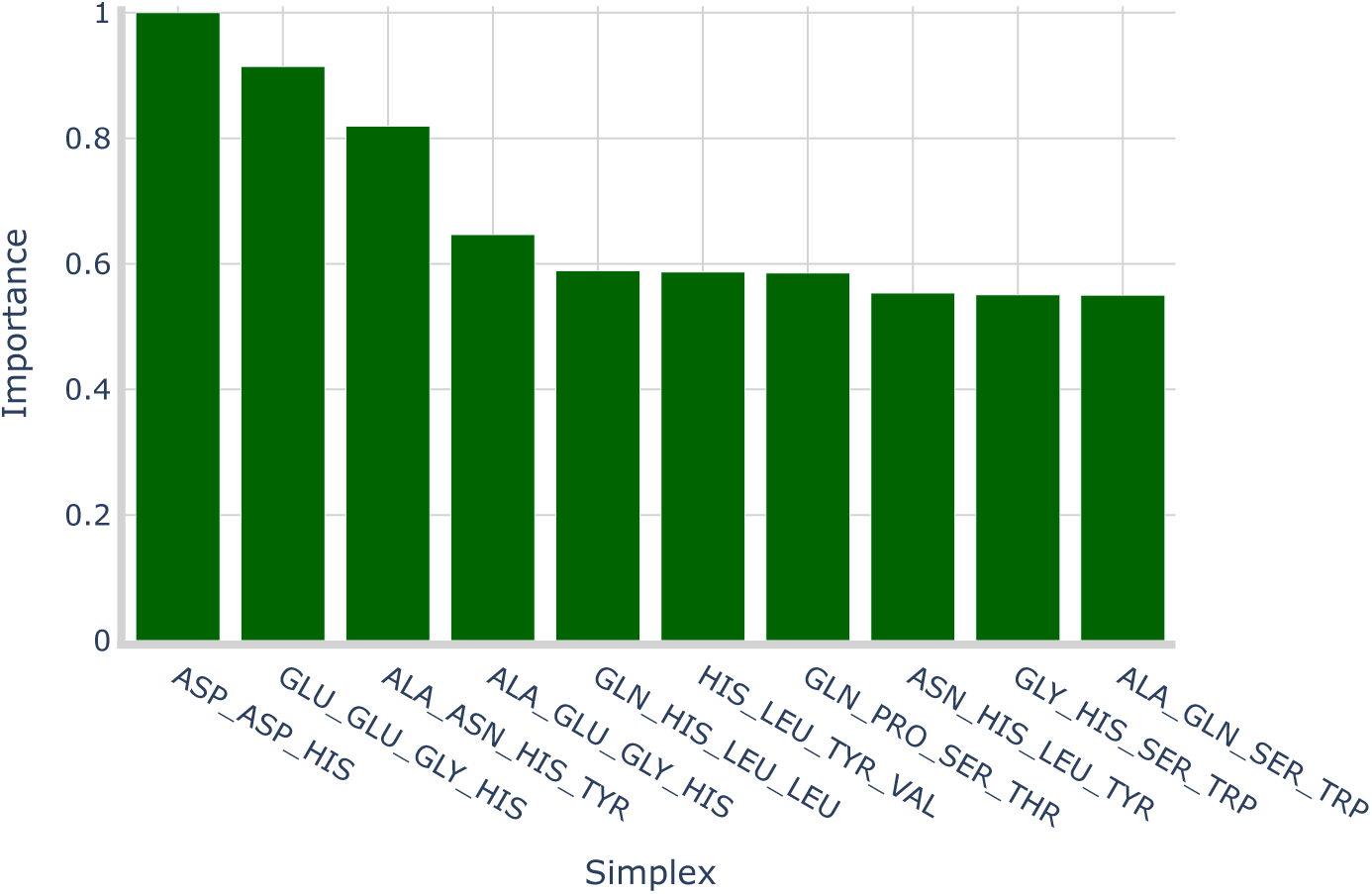
Top 10 most relevant simplices according to RF (MDI) on the INDVAL-filtered embedding – Task B.

**Figure 13:**
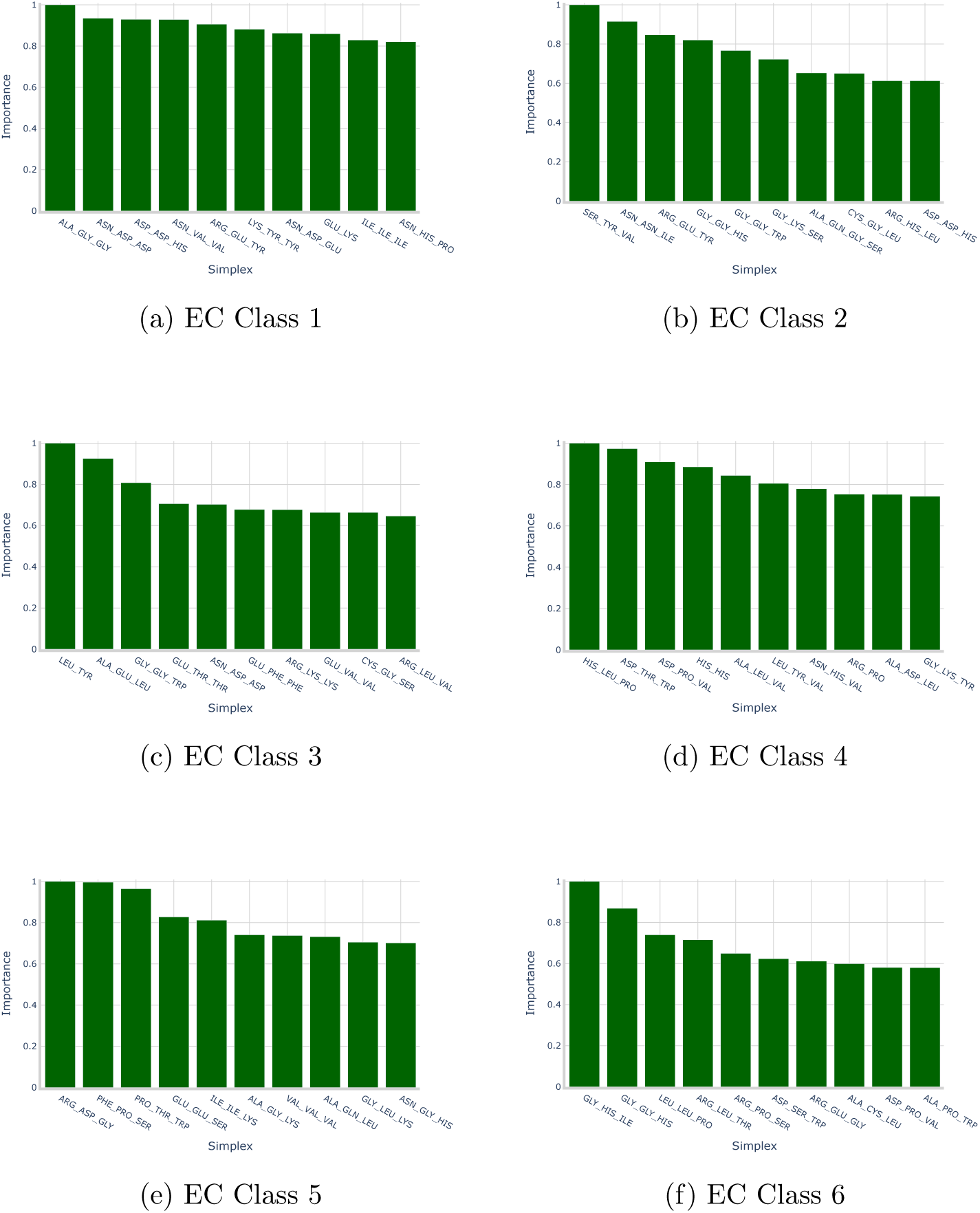
Top 10 most relevant simplices for each EC class according to *ℓ*_1_-SVM (coefficients) on the INDVAL-filtered embedding – Task B. such situations.

#### 6.2.3 Hypergraph Kernels

Table 17 shows the *ν*-SVM test performances on both kernel methods in Task B. In the multiclass setting of Task B, the kernel-based approaches remain competitive but exhibit a reversed ranking with respect to Task A: the HCK attains an ABA of 0.898, surpassing the WJK with ABA of 0.884. This pattern suggests that, when discriminating among enzyme classes, cosine similarity over sparse symbolic histograms may be more tolerant to inter-class sharing of substructures, whereas Jaccard’s overlap ratio can over-penalize

**Table 17:** Test set performances with kernel methods – Task B.

| Model | Kernel | ABA | ACC | F1 |
| --- | --- | --- | --- | --- |
| ν-SVM | HCK | $0.898 \pm 0.011$ | $0.962 \pm 0.004$ | $0.962 \pm 0.004$ |
| | WJK | $0.884 \pm 0.007$ | $0.963 \pm 0.003$ | $0.963 \pm 0.003$ |

#### 6.2.4 Spectral Density Embedding

Table 18 presents the testing performances for all models combined with Spectral Embedding in Task B. Compared to the Task A scenario the *ν*-SVM model clearly outperforms the other two candidates and confirms itself as the best model for enzyme classification based on spectral density also in a multiclass setting.

**Table 18:** Test set performances on spectral embedding – Task B.

| Model | ABA | ACC | F1 |
| --- | --- | --- | --- |
| ν-SVM | $0.722 \pm 0.015$ | $0.854 \pm 0.009$ | $0.853 \pm 0.009$ |
| $\ell_1$-SVM | $0.424 \pm 0.009$ | $0.567 \pm 0.006$ | $0.563 \pm 0.006$ |
| RF | $0.682 \pm 0.012$ | $0.805 \pm 0.008$ | $0.806 \pm 0.007$ |

The *ℓ*_1_-SVM model performances are stained by high linear correlation in the input dataset also in the multiclass scenario. This behaviour is expected, as the correlation structure of the X*_A_*^(SPECTRAL)^ is virtually identical to that of X*_B_*^(SPECTRAL)^, given that the former is obtained by simply filtering the latter to retain only enzymatic proteins, without altering the underlying feature construction.

#### 6.2.5 Multiclass GNN

Table 19 presents GNN testing performances for Task B. The same training pattern presented in Section 6.1.5 was used resulting in a different topology for each of the data splits. The trends encountered for Task B are similar to the ones of Task A: 4 out of 5 of the topologies exploit OHE representation of amino acid labels and 3 out of 5 of them chose GIN Convolution as MP strategy. Coherently with Task A, 4 out of 5 of the structures used max pooling and all of them exploited batch normalization in MP layers. The biggest differences with respect to the results of Task A in terms of GNN topologies are the increased number of hidden dimensions (4 out of 5 of the topologies leveraged the maximal available value of 256) and the ubiquitous presence of normalization layers after pooling. The sharp increase in hidden dimensions suggests that the multiclassification setting of Task B represents an inherently more complicated task, in this context more flexibility and expressive power (e.g., a wider GNN structures) bring better performances without incurring in overfitting and degraded test performances.

**Table 19:** GNN test set performances – Task B.

| **Model** | **ABA** | **ACC** | **F1** |
| --- | --- | --- | --- |
| GNN | $0.921 \pm 0.011$ | $0.942 \pm 0.008$ | $0.946 \pm 0.006$ |

#### 6.2.6 Summary of Task B

In Task B, multiclass enzyme classification benefits markedly from representations grounded in higher-order structural motifs. As summarized in Table 20, spectral density embeddings are consistently the weakest option across models, mirroring the limitations observed in Task A. In contrast, both the simplicial complexes and INDVAL embeddings achieve substantially higher ABA, with *ℓ*_1_-SVM emerging as the most effective classifier for explicit histogram representations. The superiority of *ℓ*_1_-SVM over *ν*-SVM and RF in this setting corroborates the advantage of embedded feature selection in very high-dimensional, sparse spaces: by shrinking most coefficients to zero, the model isolates a compact set of class-discriminative simplices and delivers optimal performances.

**Table 20:** Test set ABA for all models and representation strategies – Task B.

| Representation Strategy | ν-SVM | $\ell_1$-SVM | RF | GNN |
| --- | --- | --- | --- | --- |
| **Spectral Density** | $0.722 \pm 0.015$ | $0.424 \pm 0.009$ | $0.682 \pm 0.012$ | - |
| **Simplicial Complexes** | $0.864 \pm 0.009$ | $0.902 \pm 0.013$ | $0.882 \pm 0.009$ | - |
| **INDVAL** | $0.861 \pm 0.008$ | $0.893 \pm 0.012$ | $0.888 \pm 0.010$ | - |
| **Histogram Kernel** | $0.898 \pm 0.011$ | - | - | - |
| **Jaccard Kernel** | $0.884 \pm 0.007$ | - | - | - |
| **Raw PCNs** | - | - | - | **$0.921 \pm 0.011$** |

Kernel approaches provide a competitive alternative without explicit feature engineering. However, the ranking between kernels reverses relative to Task A: the HCK surpasses the WJK in ABA terms both in validation and testing.

Finally end-to-end graph neural networks trained on raw PCNs deliver the strongest overall results in Task B. The optimized GNNs reach a test ABA of 0.921. The prevalence of wider hidden dimensions, post-pooling normalization and MP operators with higher representational capacity (e.g., GIN Convolution) indicate that multiclass EC prediction benefits from increased model expressivity.

The difference between the GNN and the second-best pipeline, the *ℓ*_1_-SVM trained on the simplicial complexes embedding, was further evaluated using McNemar’s test. A statistically significant difference in favor of the GNN was observed in three of the five test splits, whereas the null hypothesis of equal error rates could not be rejected in the remaining two. These results provide evidence that the GNN generally outperforms the competing pipeline in Task B, although the advantage was not statistically consistent across all data splits.

These findings combined show that first-level EC assignment can be addressed effectively by both classical machine learning and deep learning approaches; among explicit representations, simplices-based histograms with *ℓ*_1_-regularized linear classification offer an attractive accuracy-efficiency-interpretability trade-off, while end-to-end GNNs constitute the optimal choice in terms of pure predictive performance with minimal feature engineering.

Table 21 shows the per-class average results of GNNs in Task B. They achieve consistently high values across most evaluation metrics, particularly for the well-represented classes EC 2 and EC 3, where both recall and precision remain above 0.95 on average. These outcomes indicate that the classifier is highly effective at capturing the structural signatures of the dominant enzyme families. Conversely, the least represented classes (EC 5 and EC 6) exhibit lower stability, with EC 6 in particular showing reduced precision and F1-score, reflecting the greater difficulty of learning from small sample sizes. Nonetheless, the model demonstrates remarkable specificity across all classes, exceeding 0.97 in every case and reaching 0.998 for EC 4, highlighting its robustness in avoiding false positives. Overall, the model not only generalizes well on abundant classes but also maintains strong discriminative power across the class spectrum despite substantial data imbalance.

**Table 21:** GNN per-class performance metrics on the test set – Task B.

| EC Class | Recall | Precision | F1-score | Specificity | Support |
| --- | --- | --- | --- | --- | --- |
| EC 1 | $0.951 \pm 0.019$ | $0.952 \pm 0.013$ | $0.952 \pm 0.014$ | $0.992 \pm 0.002$ | 591 |
| EC 2 | $0.928 \pm 0.010$ | $0.973 \pm 0.006$ | $0.950 \pm 0.004$ | $0.982 \pm 0.004$ | 1776 |
| EC 3 | $0.954 \pm 0.009$ | $0.956 \pm 0.008$ | $0.955 \pm 0.007$ | $0.978 \pm 0.004$ | 1437 |
| EC 4 | $0.975 \pm 0.009$ | $0.978 \pm 0.008$ | $0.977 \pm 0.007$ | $0.998 \pm 0.001$ | 311 |
| EC 5 | $0.942 \pm 0.013$ | $0.892 \pm 0.027$ | $0.916 \pm 0.013$ | $0.996 \pm 0.001$ | 132 |
| EC 6 | $0.854 \pm 0.053$ | $0.460 \pm 0.059$ | $0.594 \pm 0.045$ | $0.978 \pm 0.007$ | 89 |

## 7 Discussion and Conclusions

This work investigated whether protein structures represented as PCNs carry sufficient topological signal to support accurate prediction of physiological roles. Two complementary classification settings were addressed on a cu-rated subset of the human proteome: a binary discrimination between enzymatic and non-enzymatic proteins (Task A) and a multiclass assignment of first-level EC classes for enzymatic proteins (Task B). The analysis involved 48, 019 proteins for Task A and 21, 679 enzymes for Task B, with all experiments conducted under a stratified data splitting protocol using fixed splits across methods to enable a systematic paired comparison between representation strategies and learning algorithms.

Both explicit graph embeddings and end-to-end graph learning were considered for the experiments. Within explicit representations, three families were explored: (i) spectral density descriptors of the normalized Laplacian associated with PCNs; (ii) symbolic histograms derived from simplicial complexes evaluated on clique hypergraphs of PCNs; and (iii) their INDVAL-filtered variants for model-agnostic feature reduction. In parallel, two graph kernels over the complete simplicial complexes embedding (the HCK and the WJK) were employed to capture non-linear similarities without additional feature engineering. Lastly, MP GNNs, were optimized to learn directly from residue–residue contact graphs in an end-to-end fashion.

Three standard classifiers (kernelised *ν*-SVM, *ℓ*_1_-SVM, and RF) were selected to probe complementary learning strategies: maximum-margin learning with a non-linear kernel, embedded feature selection via *ℓ*_1_ regularization, and non-parametric ensembles sensitive to feature interactions. End-to-end GNNs on the other hand were tuned over a plethora of architectural choices to identify effective PCN encoders under feasible computational constraints. Results were satisfactory for both tasks: in Task A, the top-performing pipeline was *ν*-SVM on WJK with test ABA of 0.900, marginally ahead of the GNN counterpart (ABA = 0.898). This phenomenon indicates that end-to-end learning can almost match kernel baselines without additional hand-crafted features.

The results from Task B strengthen such considerations as end-to-end GNNs were the best overall with test ABA = 0.921, while among explicit representations the *ℓ*_1_-SVM on the full simplicial histogram reached the best results with test ABA = 0.902. Per-class analysis of GNN results showed excellent precision and recall on well-represented classes (EC classes 1–4) with slight instability on sparser classes (EC classes 5–6).

An additional insight emerges from the consistently lower performance of the spectral density representation across both tasks. As the spectral density is the only tested representation that provides a compact summary of the global connectivity organization of a PCN without explicitly considering its local substructures or residue labels, its weaker predictive performance suggests that physiological role information is more readily captured through localized residue-level patterns than through global summaries alone.

The consistent emergence of the asp-asp-his 2-simplex as a top discriminant is coherent with well-established enzymological knowledge and provides a biological rationale for its predictive value in both Task A and Task B. Histidine–aspartate interactions constitute one of the most recurrent catalytic building blocks in enzyme active sites: the hydrogen bond between histidine and aspartate is central to the classical ser–his–asp catalytic triad of serine proteases, where the aspartate tunes the p*K*_a_ of the histidine and enables it to act as a general base, with mutation of the triad aspartate reducing catalytic efficiency by up to four orders of magnitude [15, 66]. Analogous his–asp dyads have been characterised in ribonuclease A [62], in the human DNA-repair endonuclease Ape1, where the pair stabilises the pentavalent transition state of phospho-transfer [40], and in glycoside hydrolases, where a his–asp pair operates as a general-acid dyad through low-barrier hydrogen bonding [59]. More broadly, histidine and aspartate/glutamate residues jointly coordinate the metal cofactor of non-heme iron dioxygenases within the strictly conserved 2-His–1-carboxylate facial triad [26], and this type of geometric arrangement recurs across otherwise unrelated folds and reaction mechanisms. The presence of two aspartates near a histidine, as captured by the asp–asp–his simplex, is thus consistent with a spatial reinforcement of a catalytic or metal-coordinating his–asp unit by a second, geometrically proximal carboxylate residue, either contributing to p*K*_a_ modulation of the histidine or participating directly in substrate or cofactor binding. Because the simplex is constructed from spatial contacts rather than sequence adjacency, it captures three-dimensional co-localisation patterns that are hardly identifiable through linear sequence motifs, which may in part explain its strong discriminative value in distinguishing enzymatic from non-enzymatic proteins and, to a lesser extent, in separating first-level EC classes.

Nonetheless, several limitations should be acknowledged: (i) the dataset excluded proteins with multiple first-level EC annotations and therefore did not model functional multiplicity; the complementary multi-label setting was investigated separately in a companion paper specifically devoted to multifunctional proteins [11]; (ii) the study focused exclusively on human proteins, and cross-species generalization was therefore not considered.

Future studies could extend end-to-end architectures to 3D-aware, E(3)-equivariant GNNs and geometric vector perceptrons to better exploit fine-grained geometric information (deliberately discarded in this work as per Section 2). Building on the complementary multi-label analysis of multifunctional proteins presented in [11], subsequent work could investigate broader formulations of functional multiplicity and annotation, including finer-grained hierarchical EC annotations. Furthermore, deep topological learning strategies—such as message passing on simplicial or cell complexes with specialized operators—could be incorporated to encode higher-order interactions and global shape priors that remain inaccessible to purely pairwise GNN models.

Overall, the findings indicate that structural information encoded in PCNs is highly predictive of protein physiological roles at scale. Topology-driven embeddings provide accurate and interpretable baselines, while carefully tuned GNNs deliver the strongest multiclass performance, opening a path toward higher order, geometry-aware, and explicitly multi-label structural analysis.

^1^https://www.rcsb.org/

